# Learning to read a second language establishes a parallel L2 representation alongside the native one in the VWFA

**DOI:** 10.64898/2026.07.16.739038

**Authors:** Siyi Fan, Xiaoxia Feng, Xi Yu, Minye Zhan, Manli Zhang, Guosheng Ding, Xiangzhi Meng

## Abstract

Learning to read a second language requires the brain to incorporate a new writing system into an already established native-language reading network, yet how this process reshapes the Visual Word Form Area (VWFA) remains poorly understood. Using fMRI with a passive viewing paradigm, we systematically investigated how VWFA responses to Chinese (L1) and English (L2) words evolved across three groups of native Mandarin-speaking children at distinct stages of L2 literacy acquisition: L2 pre-readers, L2 beginning readers, and L2 advanced readers. We combined univariate activation analyses, representational similarity analysis, and supervised machine learning classification to investigate two theoretical accounts of VWFA reorganization: the Overlay Model, which predicts stable L1 responses as L2 responses emerge, and the Competing Model, which predicts competitive reallocation of neural resources from L1 to L2. We found that although the VWFA already exhibited robust selective responses to L1 words, L2-word selectivity was absent in L2 pre-readers but emerged robustly in beginning readers, with L1-word selectivity remaining stable throughout. Representational similarity analysis further revealed that robust L2 word representations within the VWFA emerged only after children began learning to read L2, while L1 word representations remained stable across all three groups. Finally, supervised machine learning analyses successfully discriminated among the three L2 literacy groups on the basis of L2 but not L1 activation patterns, indicating that VWFA responses to L2 alone were sufficient to capture children’s stages of L2 literacy acquisition, whereas responses to L1 were insensitive to L2 learning experience, providing no support for the Competing Model’s prediction of competitive neural reallocation away from L1. Together, these findings support the Overlay Model, demonstrating that the VWFA incorporates a new writing system within its existing cortical resources without compromising L1 print processing.

## 1 Introduction

Learning to read is a cognitively demanding process that entails years of practice and formal instruction, marking a pivotal milestone in children’s cognitive development. Despite growing interest in how learning to read reorganizes the human brain, fMRI studies in pediatric populations, particularly in pre-readers, remain scarce and have focused predominantly on the acquisition of a single native writing system (Brem et al., 2010; Chyl et al., 2018; Dębska et al., 2024; Dehaene-Lambertz et al., 2018; Yeatman et al., 2024; Yu et al., 2018). Existing studies have nevertheless provided robust evidence that the Visual Word Form Area (VWFA) is a hallmark of the literate brain, as it gradually emerges and becomes increasingly specialized for print processing in both children (Centanni et al., 2018; Feng et al., 2020; Kubota et al., 2019; Monzalvo et al., 2012) and previously illiterate adults (Hervais-Adelman et al., 2019). Yet increasing numbers of children worldwide are becoming bilingual readers, most commonly as sequential bilinguals who acquire second-language (L2) literacy after establishing an initial foundation in their native language (L1). Despite this reality, the neural basis of bilingual literacy acquisition remains poorly understood, particularly how a reading system already tuned to the native writing system incorporates a second writing system.

Prior research in adult bilinguals has examined how the VWFA processes two writing systems, revealing various neural representation patterns across different scripts (Bai et al., 2011; Baker et al., 2007; Nelson et al., 2009; Wong et al., 2009; Xu et al., 2017; Zhan et al., 2023). Specifically, while some studies reported substantial spatial overlap in the VWFA for both scripts, as seen in English–Hebrew, Chinese–Korean, and Chinese–English bilinguals (Bai et al., 2011; Baker et al., 2007; Wong et al., 2009), others employing multivariate pattern analyses or high-resolution 7T fMRI have unveiled script-specific internal representations and partially segregated cortical patches (Xu et al., 2017; Zhan et al., 2023). Crucially, because these findings were derived from fluent adult bilinguals, they merely reflect the stable end-state of years of reading experience. A more fundamental and still unresolved question, therefore, is how the cortical territory in the ventral visual cortex, encompassing the VWFA, initially reorganizes when a child first begins to acquire a second writing system.

In parallel, extensive research has investigated how word-selective responses emerge in the VWFA during native-script reading acquisition, motivating several theoretical accounts (Dehaene & Cohen, 2007; Dehaene-Lambertz et al., 2018; E. Kubota et al., 2024). The neuronal recycling hypothesis (Dehaene & Cohen, 2007) proposes that neural populations originally tuned to other visual categories are progressively co-opted for orthographic processing, implying a competition between words and other visual representations. In support of this view, higher reading ability in illiterate and literate adults was associated with a modest reduction in face responsivity in the left fusiform gyrus, accompanied by a corresponding rightward shift in the hemispheric lateralization of face processing (Dehaene et al., 2010). Similarly, in 6-year-old children with varying reading performance, the spatial extent of letter-selective regions (VWFA) was negatively correlated with the extent of the left fusiform face area (FFA) (Centanni et al., 2018). This competitive account explains clinical patterns as well: individuals with dyslexia exhibit reduced word selectivity in the VWFA, which was driven by elevated responses to objects rather than weaker responses to words (Kubota et al., 2019). Moreover, recent longitudinal studies have found that, over development, increases in word selectivity were directly linked to decreases in limb selectivity, rather than face or object selectivity (Nordt et al., 2021, 2023). These findings leave the competitive mechanism underlying VWFA development unclear.

However, some studies challenge this competitive account, reporting that the emergence of word-selective responses was not accompanied by suppression of other visual categories (Dehaene-Lambertz et al., 2018; Feng et al., 2022; Hervais-Adelman et al., 2019). In our previous work, we empirically investigated how the VWFA emerged during native-script reading acquisition through a cross-sectional design comprising three groups of children: 6-year-old pre-readers, 6-year-old beginning readers, and 9-year-old advanced readers (Feng et al., 2022). We found that word-selective responses in the VWFA were absent in pre-readers but emerged robustly in beginning readers of the same age, confirming that VWFA emergence is driven by reading experience rather than maturation or age. Critically, the emergence of word selectivity in the VWFA reflected increased activation to written words rather than decreased responses to other visual categories, arguing against the strict cortical encroachment posited by the original neural recycling hypothesis derived from adult illiterates (Dehaene et al., 2010). This pattern of findings is consistent with the revised neuronal recycling account proposed by Dehaene-Lambertz et al. (2018), and is further supported by adult literacy evidence from Hervais-Adelman et al. (2019), suggesting that VWFA emergence may occur without necessarily eliminating pre-existing responses to other visual categories.

Importantly, whether the mechanisms underlying VWFA emergence extend to L2 literacy acquisition in children who have already developed native-script selectivity in the VWFA remains an open question. While our prior work demonstrated that learning a native script does not encroach upon the cortical territory of non-text categories like faces, it remains entirely unknown whether this non-competitive principle also holds within the orthographic domain. To characterize the nature of this cortical reorganization induced by L2 literacy acquisition, we propose two potential theoretical accounts: the Overlay Model and the Competing Model. The Overlay Model posits that L2 literacy is acquired within the ventral visual cortex as a functional “overlay,” without imposing neural interference on L1 print processing. The Competing Model, conversely, holds that L2 acquisition triggers competition that reallocates neural resources away from L1, thereby compromising its pre-existing print processing.

In the current study, we asked three questions: (1) whether L2 literacy acquisition induces neurofunctional reorganization in the ventral visual cortex which encompasses the VWFA already selective for the native writing system; (2) if so, where L2-selective responses emerge, specifically whether they are spatially aligned with pre-existing L1-selective responses or instead exhibit a systematic spatial shift; and (3) how they emerge, that is, whether L2 acquisition proceeds as a functional overlay leaving L1 intact, or as a competitive process that disrupts pre-existing L1 representation, thereby adjudicating between the Overlay and Competing Models. To address these questions, we focused on Chinese–English bilingual acquisition, as Chinese and English represent a maximally contrastive pairing of writing systems: Chinese is a morphosyllabic logographic script characterized by complex spatial configurations, whereas English employs an alphabetic system with relatively transparent grapheme-to-phoneme correspondences. This structural distance makes Chinese–English an ideal case for probing the neural mechanisms underlying bilingual literacy acquisition in the developing brain.

We recruited native Mandarin-speaking children across three stages of English literacy development: L2 pre-readers, beginning readers, and advanced readers. These three groups were age-matched but differed critically in their English literacy experience, with the L2 pre-reader group having received no or limited formal English literacy training, providing a rare opportunity to isolate the contribution of L2 literacy experience from maturational factors. As in our previously published fMRI studies of reading (Feng et al., 2020, 2022), all children performed the same passive viewing task with words, faces, and houses, with the sole task of detecting an occasional target star. To address the above three questions, we combined complementary univariate, multivariate, and machine-learning analyses. In the univariate analyses, we first examined category-selective activation across and within each group at both the whole-brain level and the ROI level. For the latter, we employed a moving-window approach along the lateral–medial and posterior–anterior gradients centered on the VWFA to clarify whether L2-selective responses emerged spatially at the same cortical territory as L1-selective responses. To ensure that group-level patterns reflected consistent individual profiles rather than averaging artifacts, we conducted individual-based voxel-wise analyses comparing the most responsive voxels for English and Chinese words within each child. Furthermore, multivariate pattern analysis was applied to test how stable L2 representations emerged and whether their emergence was accompanied by disruption of L1 representations. Finally, we utilized supervised machine learning classification to examine whether neural patterns evoked by L2 and L1 words could predict L2 literacy stage at the individual level, and critically, whether successful classification would be restricted to L2-based neural patterns or would also extend to L1-based neural patterns. Classification based on L1 patterns would indicate that L1 representations are being reshaped by L2 acquisition, as specifically predicted by the Competing Model. Together, this design allowed us to characterize how L2 literacy experience shapes neural responses to a newly learned writing system and directly test the Overlay and Competing Models, thereby clarifying the neural mechanisms by which the developing brain accommodates a typologically distinct second writing system.

## 2 Methods

### 2.1 Participants

The present study initially recruited 60 typically developing children, all of whom were native Chinese speakers and primary school students. Despite being of similar age, participants varied substantially in their L2 literacy experience and were therefore assigned to three groups: L2 pre-readers, L2 beginning readers, and L2 advanced readers. The observed differences in L2 reading performance among children of the same age can be partly attributed to the educational context in China, where English is formally introduced as a second language starting in the third grade. However, some children had exposure to English before formal schooling through interest-based activities or extracurricular tutoring, resulting in substantial individual differences in early English literacy. Two participants were excluded from subsequent analyses because of excessive head motion during fMRI scanning. The final sample thus comprised 58 children with high-quality fMRI images (mean age = 10.18 ± 1.04 years; 27 females, 31 males), including 20 L2 pre-readers, 19 L2 beginning readers, and 19 L2 advanced readers. A sensitivity power analysis in G*Power 3.1 showed that the final sample size of 58 participants provided 80% power to detect medium-to-large group effects in a one-way ANOVA with three groups at α = .05 (minimum detectable effect size: *f* = .42, η_p_² ≈ .15). None of the children in the current study had a history of neurological, psychiatric, or hearing impairments. Written informed consent was obtained from both children and their parents. This study received approval from the Research Ethics Committee of Beijing Normal University.

Children’s L2 reading performance was assessed using a battery of English literacy tests, including a lab-developed spelling test (Meng et al., 2016; You et al., 2011) and the Word Identification and Word Attack subtests from the Woodcock Reading Mastery Test (Woodcock, 1998). In the Spelling test (total score = 40), children were required to write down English letters/words upon their auditory presentation (Meng et al., 2016; You et al., 2011). The Word Identification test (total score = 106) measures children’s ability to read isolated words aloud, while the Word Attack test (total score = 45) assesses the skill in applying phonics and structural analysis skills to decode unfamiliar pseudo-words. Additionally, principal component analysis (PCA) was performed based on these three tests to derive a composite score reflecting each child’s L2 literacy proficiency. This composite score was subsequently entered as a predictor in the brain–behavior regression analyses.

The L2 pre-reader group comprised 20 children (mean age = 9.84 ± 0.97 years; 6 females, 14 males). Their pre-reader status was determined based on a combination of three aspects of early English literacy: limited single-word reading ability (assessed via Word Identification), limited phonological lexical representation (assessed via Spelling), and a lack of grapheme-phoneme correspondence knowledge (assessed via Word Attack). Specifically, children in the pre-reader group showed lower performance across the three English tests, with raw scores ranging from 9 to 18 on the Word Identification test, 1 to 8 on the Spelling test, and 0 to 7 on the Word Attack test. Although some children in the pre-reader group achieved relatively higher scores on Word Identification, this performance may have resulted from incidental exposure or rote memorization. Their spelling and nonword decoding performance nevertheless remained very limited, indicating that they had not yet developed stable orthographic representations and grapheme–phoneme correspondence knowledge. Overall, this profile closely resembled that of the L1 pre-readers in our previous study, who read fewer than 15 words per minute (Feng et al., 2022).

In comparison, the beginning reader group consisted of 19 children (mean age = 10.19 ± 1.12 years; 11 females, 8 males) who were comparable in age to the pre-readers but demonstrated greater L2 literacy experience with higher performance across the above three English tests (Table 1). Finally, the advanced reader group consisted of 19 children (mean age = 10.51 ± 0.97 years; 10 females, 9 males) who were also comparable in age to the pre-readers and beginning readers but had substantially greater L2 literacy experience and proficiency **(Table 1)**. This group served as a higher-literacy comparison group for examining how increasing levels of L2 literacy experience and proficiency are associated with brain organization.

**Table 1.**
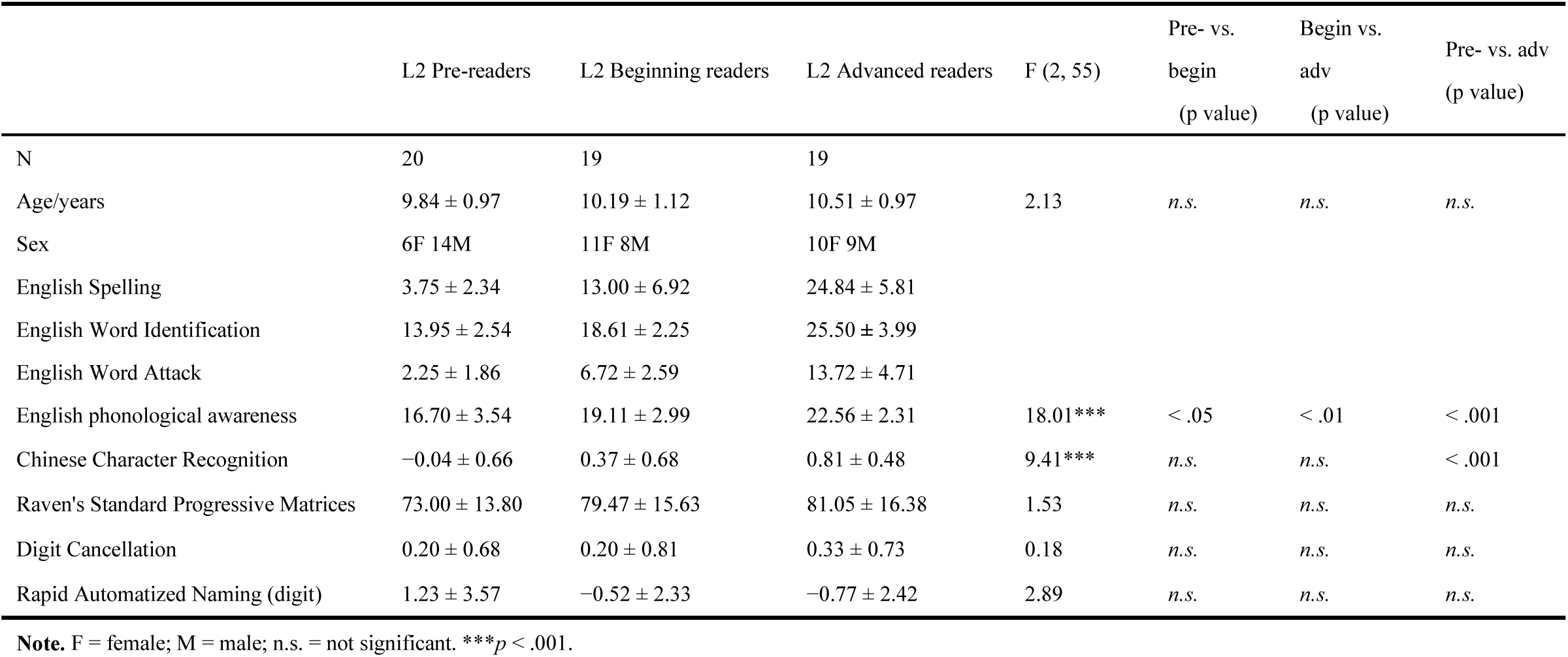
Characteristics of the three groups (means and standard deviations).

|  | L2 Pre-readers | L2 Beginning readers | L2 Advanced readers | F (2, 55) | Pre- vs.<br>begin<br>(p value) | Begin vs.<br>adv<br>(p value) | Pre- vs. adv<br>(p value) |
| --- | --- | --- | --- | --- | --- | --- | --- |
| N | 20 | 19 | 19 |  |  |  |  |
| Age/years | 9.84 ± 0.97 | 10.19 ± 1.12 | 10.51 ± 0.97 | 2.13 | *n.s.* | *n.s.* | *n.s.* |
| Sex | 6F 14M | 11F 8M | 10F 9M |  |  |  |  |
| English Spelling | 3.75 ± 2.34 | 13.00 ± 6.92 | 24.84 ± 5.81 |  |  |  |  |
| English Word Identification | 13.95 ± 2.54 | 18.61 ± 2.25 | 25.50 ± 3.99 |  |  |  |  |
| English Word Attack | 2.25 ± 1.86 | 6.72 ± 2.59 | 13.72 ± 4.71 |  |  |  |  |
| English phonological awareness | 16.70 ± 3.54 | 19.11 ± 2.99 | 22.56 ± 2.31 | 18.01*** | < .05 | < .01 | < .001 |
| Chinese Character Recognition | -0.04 ± 0.66 | 0.37 ± 0.68 | 0.81 ± 0.48 | 9.41*** | *n.s.* | *n.s.* | < .001 |
| Raven's Standard Progressive Matrices | 73.00 ± 13.80 | 79.47 ± 15.63 | 81.05 ± 16.38 | 1.53 | *n.s.* | *n.s.* | *n.s.* |
| Digit Cancellation | 0.20 ± 0.68 | 0.20 ± 0.81 | 0.33 ± 0.73 | 0.18 | *n.s.* | *n.s.* | *n.s.* |
| Rapid Automatized Naming (digit) | 1.23 ± 3.57 | -0.52 ± 2.33 | -0.77 ± 2.42 | 2.89 | *n.s.* | *n.s.* | *n.s.* |

**Note.** F = female; M = male; *n.s.* = not significant. \*\*\* $p < .001$ .

Descriptive statistics for English Spelling, Word Identification, and Word Attack are presented in Table 1, as these measures were used to characterize children’s L2 literacy status. The three groups also differed in English phonological awareness, an English-related measure that was not used for group classification (*F*(2, 55) = 18.01, *p* < .001, η² = .41). Age did not differ significantly across groups (*p* > .05). To evaluate whether group differences in L2 literacy were confounded by L1 literacy ability, Chinese character recognition was also assessed and was found to differ across groups (*F*(2, 55) = 9.41, *p* < .001, η² = .26), with only pre-readers scoring lower than advanced readers. To examine whether lower L2 literacy performance in pre-readers was associated with individual differences in L1 literacy ability, we examined within-group correlations between Chinese character recognition and English literacy measures. These correlations were not significant in the L2 pre-readers or beginning readers (*ps* > .05), whereas among advanced readers, Chinese character recognition was significantly correlated with both English Spelling and English Word Identification (**Supplementary Table S1**).

The children also completed several cognitive assessments: Raven’s Standard Progressive Matrices to measure nonverbal reasoning (Zhang & Wang, 1989), Digit Cancellation test (Mirsky et al., 1991) to assess sustained attention and rule out attention deficits, and a digit-based Rapid Automatized Naming task (Pan et al., 2011) to measure rapid naming ability. No significant group differences were found in Raven’s scores, digit cancellation, or digit RAN performance (*ps* > .05; Table 1).

### 2.2 Stimuli and task

The stimuli and experimental paradigm were adapted from our previous studies (Feng et al., 2020, 2022). During MRI scanning, five categories of visual stimuli (English words, Chinese words, faces, houses and rotating checkerboard) were presented in a mini-block design. The stimulus set comprised 30 four-letter English words, 30 single-character Chinese words, 30 non-famous Asian faces (frontal or slightly angled), and 30 house images. All stimuli were gray-scale, presented in black on a white background, with luminance and image size matched across conditions.

All children completed two functional runs. Each run consisted of nine mini-blocks presented in randomized order: two blocks each of English words, Chinese words, faces, and houses, and one block of checkerboard. Each mini-block lasted 18 s and was followed by a 10.5 s fixation cross. In each block, 10 pairs of different images belonging to the same category (200 ms presentation for the first picture/word, 200 ms inter-stimulus interval, 500 ms presentation for the second picture/word) were presented, separated by a 600 ms fixation period. In addition to the 10 image pairs, each block contained two target stars (1500 ms each) presented at random positions. Children were instructed to press a button with their right index finger whenever a star appeared. This task was designed to keep the children’s attention focused on the visual stimuli, but without any explicit word-reading requirement.

Before scanning, all participants completed a brief training session in a mock MRI scanner to familiarize them with the scanning environment, task procedure, and the requirement to minimize head movement.

### 2.3 fMRI data acquisition

All MRI data were collected on a Siemens Tim Trio 3T scanner at the Beijing Normal University’s imaging center. Stimuli were presented using E-Prime 2.0, and children viewed them through a mirror mounted on the head coil while wearing MRI-compatible noise-attenuating earphones. Functional images were acquired using a 12-channel head coil and a gradient-echo planar imaging (EPI) sequence with the following parameters: TR = 2400 ms, TE = 30 ms, flip angle = 81°, matrix = 64×64, voxel size = 3×3×3 mm³, slice thickness = 3 mm, number of slices = 40 (contiguous ascending acquisition). A T1-weighted volume was also acquired with the following parameters: 176 axial slices, TR = 2300 ms, TE = 4.18 ms, flip angle = 9°, matrix = 256×256.

### 2.4 fMRI data preprocessing

Preprocessing of the fMRI data was performed using DPABI (Yan et al., 2016). Images were corrected for slice timing and realigned to the first volume for head-motion correction. Head motion was quantified using framewise displacement (FD). Participants were excluded if they exhibited excessive motion, defined as a mean FD > 0.5 mm, more than 20% of volumes exceeding the 0.5-mm FD threshold, or any absolute displacement greater than 3 mm. For spatial normalization, each participant’s T1-weighted image was co-registered to the mean functional image and segmented using unified segmentation; the resulting deformation parameters were then applied to the functional images to normalize them to MNI space. Nuisance regressors (cerebrospinal fluid and the six head-motion parameters from rigid-body realignment) were removed, followed by linear detrending. Finally, normalized functional images were spatially smoothed with a 6-mm FWHM Gaussian kernel. For multivariate pattern analyses, including RSA and classification, unsmoothed normalized functional images were used to preserve fine-grained spatial patterns.

### 2.5 Statistical analyses

#### 2.5.1 Category-selective activations at the whole-brain level

Functional data were analyzed using a general linear model (GLM) at the individual level. Each stimulus category (English words, Chinese words, faces, houses and checkerboard) was modeled across the two runs, resulting in five regressors of interest. Mini-blocks were modeled as boxcar functions with their actual durations and convolved with the canonical hemodynamic response function (HRF). Six head-motion parameters from rigid-body realignment were included as nuisance regressors.

Activations were examined using two complementary types of contrasts. First, category-selective contrasts were conducted to identify category-specific responses by comparing each category against the mean response to the other visual categories: (1) English words versus the mean of faces and houses, (2) Chinese words versus the mean of faces and houses, (3) faces versus the mean of English words, Chinese words, and houses, (4) houses versus the mean of English words, Chinese words, and faces. Second, category-versus-fixation contrasts were performed to estimate the overall activation associated with each category by comparing each condition with fixation. These contrasts were subsequently used to assess activation magnitude across categories.

At the group level, functional activation maps for each category-selective contrast were examined in the entire sample (N = 58) and visualized within each of the three L2 subgroups: pre-readers (N = 20), beginning readers (N = 19), and advanced readers (N = 19). Whole-sample maps were thresholded at *p* < .001 at the voxel level and *p* < .05 FWE-corrected at the cluster level. Given the smaller sample sizes and substantial inter-individual variability in neural activation within each subgroup, subgroup activation maps were presented for descriptive purposes using an uncorrected voxel-level threshold of *p* < .001 to visualize activation patterns across different stages of L2 reading development.

To examine group differences across the three groups and the association between L2 literacy proficiency and brain activation across the entire sample, two whole-brain voxel-wise analyses were conducted. First, a one-way ANOVA was performed with L2 reading group as the between-subjects factor and individual-level English-selective contrast maps as the dependent variable. Second, a whole-brain multiple regression analysis was conducted using the English-selective contrast maps as the dependent variable. The predictor of interest was a PCA-derived English literacy composite score based on Spelling, Word Identification, and Word Attack scores, with age and Chinese character recognition ability entered as covariates. Statistical maps were thresholded at p < .001 at the voxel level and *p* < .05 FWE-corrected at the cluster level. Given the stringent nature of this correction, we additionally report an exploratory analysis at two liberal thresholds (voxel-level *p* < .001 and *p* < .005, uncorrected at the cluster level; see **Supplementary Fig. S1**), which revealed voxels showing group differences and voxels associated with English literacy proficiency.

Given our strong prior hypothesis regarding the role of the VWFA in reading acquisition, we next focused on the left ventral occipitotemporal cortex (vOTC) encompassing the VWFA to investigate: (1) where L2-selective responses emerged, specifically whether they were spatially aligned with or systematically shifted from pre-existing L1-selective responses; and (2) how L2-selective responses emerged, specifically whether their emergence was accompanied by stable L1 responses or by disruption of L1 responses.

#### 2.5.2 Moving-window ROI analyses along the lateral–medial and posterior–anterior axes centered on the VWFA

To investigate the functional organization of the left vOTC and its preferential responses to different visual categories, with a focus on the spatial relationship (dispersion or alignment) of responses to L1 and L2 words, we performed gradient-based ROI analyses across three groups. For the lateral to medial gradient centered on the VWFA, β estimates for each category relative to fixation were extracted from 6-mm-radius spheres centered at four regularly spaced locations along the x-axis (x = – 57, –45, –33, –21), with the y- and z-coordinates fixed at the classical VWFA location (y = –57, z = –12). The y- and z-coordinates were determined according to previous studies on illiterates (Dehaene et al., 2010), typically developing children (Feng et al., 2020) and adults (Vogel et al., 2012). For the posterior to anterior gradient centered on the VWFA, β estimates for each category relative to fixation were extracted from 6-mm-radius spheres centered at five regularly spaced locations along the y-axis (y = – 72, –60, –48, –36, –24), with the x- and z-coordinates fixed at the classical VWFA location (x = –45, z = –12). This axis was chosen given previous evidence of a functional gradient related to orthographic and linguistic processing along the posterior-anterior axis (van der Mark et al., 2011; Vinckier et al., 2007).

For each ROI along the lateral–medial and posterior–anterior axes, the β values for each visual category relative to fixation were plotted separately for each group. These activation-profile plots aimed to objectively illustrate the gradient of categorical responses along these axes, with a primary focus on the relative spatial organization of Chinese and English word responses, especially between pre-readers and readers with L2 experience. Exploratory English–Chinese comparisons were conducted within each group at each ROI. To account for multiple ROI-wise comparisons, false discovery rate (FDR) correction was applied to the resulting *p* values.

To further quantify word-selective responses, we computed separate selectivity indices (SIs) for English and Chinese words within each ROI. This allowed us to statistically test whether L2-word selectivity emerged within the same cortical territory as L1 in the vOTC, and whether L2-word selectivity increased with L2 reading experience and approached the spatial profile of L1-word selectivity. Using the same sets of lateral–medial and posterior–anterior ROIs described above, word selectivity indices were computed as follows:

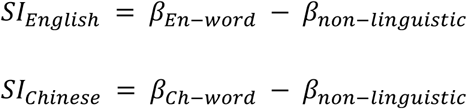

where β*_En-word_* and β*_Ch-word_* denote the extracted β estimates for English and Chinese words versus fixation, respectively, and β*_non-linguistic_* represents the mean β estimate for non-linguistic categories (faces versus fixation and houses versus fixation). A positive SI indicates a stronger response to English or Chinese words relative to non-linguistic categories, whereas a negative SI reflects greater selectivity to non-linguistic stimuli. One-way ANOVAs were then conducted for each ROI and script to examine group differences in word selectivity, followed by LSD post hoc comparisons to assess pairwise differences between the three groups.

#### 2.5.3 Individual-level analyses: Cross-run responses in the most responsive VWFA voxels

The group-level and ROI-based analyses described above characterize spatially averaged responses to two scripts and other visual categories, which assume consistent functional localization across all children. However, less proficient readers may exhibit greater anatomical variability in the spatial distribution of active brain regions, and such variability could obscure fine-grained functional differences within the VWFA even when overall activation is comparable. To test whether L2 responses emerged within the most responsive VWFA voxels without reducing pre-existing L1 responses, we conducted an individual-level voxel selection analysis within the VWFA.

For each participant, analyses were conducted within a 10-mm-radius sphere centered on the canonical VWFA coordinates (Vogel et al., 2012). Following Jacoby and Fedorenko (2018), voxels were ranked separately according to their category-selective activation for English words relative to the mean of faces and houses and for Chinese words relative to the mean of faces and houses. The top 10% of voxels were then selected for each contrast individually (**Fig. 3A**). The 10-mm-radius VWFA sphere contained 170 voxels, such that the top 10% selection comprised 17 voxels for each script. We used a two-fold cross-run selection-extraction procedure. In the first fold, Run 1 was used to define the English and Chinese top-10% voxel sets, and independent beta estimates for English and Chinese words relative to fixation were extracted from Run 2. In the second fold, Run 2 was used for voxel selection and Run 1 for beta extraction. For each participant and voxel set, the final beta estimates for English and Chinese words were obtained by averaging the extracted values across the two folds.

**Fig. 1.**
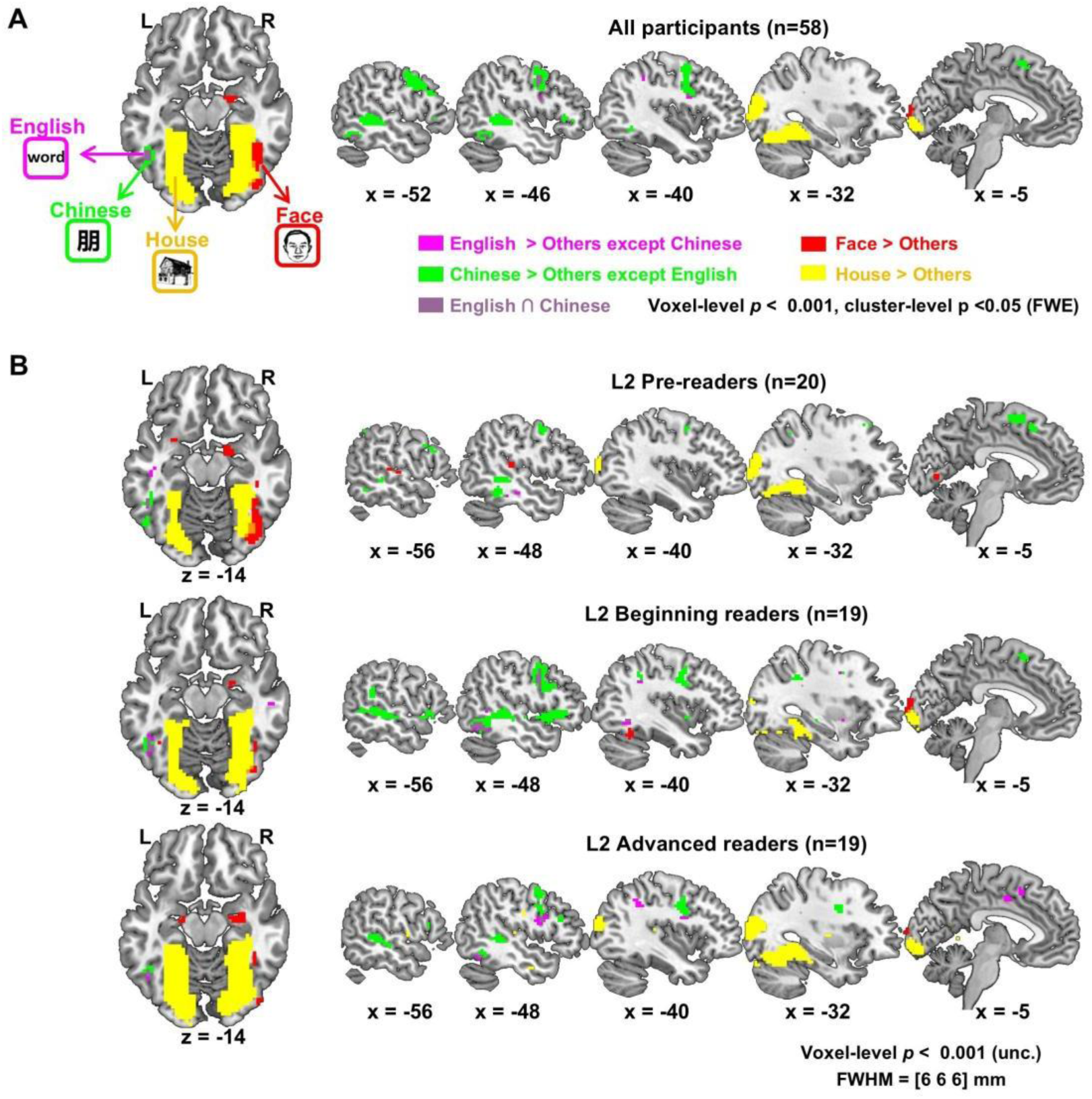
Category-selective activations. Horizontal and sagittal slices showing category-selective activations across all participants (A) and separately for L2 pre-readers (B), L2 beginning readers (C), and L2 advanced readers (D). Violet: English word-selective activations (English words > [Faces, Houses]); Green: Chinese word-selective activations (Chinese words > [Faces, Houses]); Red: face-selective activations (Faces > [Chinese words, English words, Houses]); Yellow: house-selective activations (Houses > [Chinese words, English words, Faces]).

**Fig. 2.**
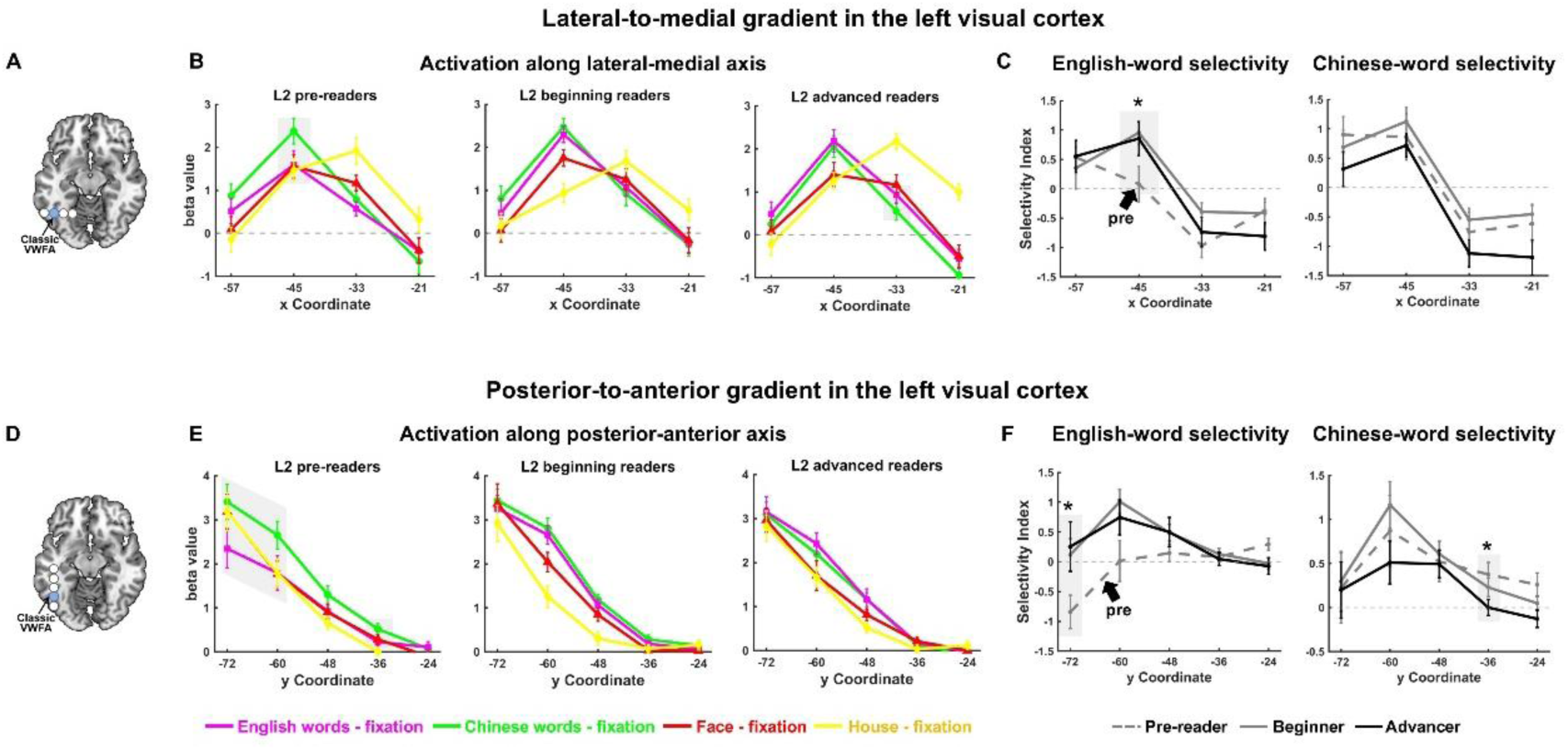
Category-selective activation gradients in the left visual cortex. (A) Lateral–medial axis in the left visual cortex examined at fixed coordinates (y = –57, z = –12) with x ranging from –57 to –21. (B) Beta values along the lateral–medial axis for English words (violet), Chinese words (green), faces (red), and houses (yellow) versus fixation for the three groups. (C) Word selectivity along the lateral–medial axis for English words (left) and Chinese words (right). Line styles denote the three groups: gray dashed for pre-readers, gray solid for beginning readers, and black solid for advanced readers. (D) Posterior-anterior axis examined at fixed coordinates (x = –45, z = –12) with y ranging from –72 to –24. (E) Beta values for each category along the posterior-anterior axis for the three groups. (F) Word selectivity along the posterior-anterior axis for English words (left) and Chinese words (right).

**Fig. 3.**
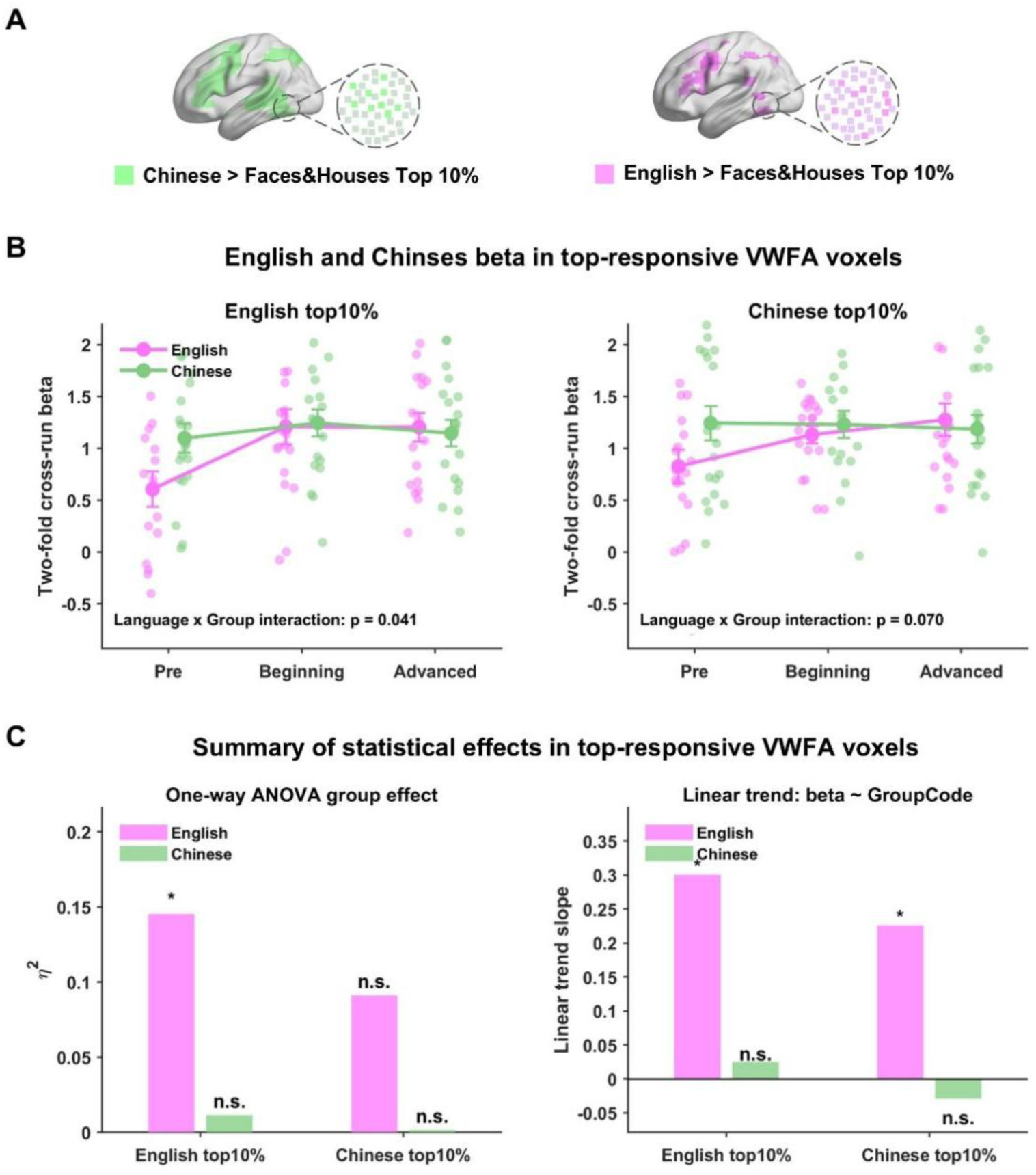
Individual-level characterization of cross-language responses within the VWFA across L2 literacy groups. (A) Voxels were ranked separately based on activation strength for the contrasts English words > the mean of faces and houses and Chinese words > the mean of faces and houses. The top 10% of voxels from each contrast were identified separately, and beta estimates were extracted using a cross-run procedure. (B) Comparison of two-fold cross-run beta estimates for English (violet) and Chinese (green) words across the three reading stages within English top-10% and Chinese top-10% voxels. (C) Summary of statistical effects, including one-way ANOVA effect sizes for group differences and linear trend slopes across the ordered L2 literacy groups, shown separately for English and Chinese responses. Significance levels: \**p* < .05; n.s., not significant.

The primary analyses focused on two individually defined voxel sets: English top-10% voxels and Chinese top-10% voxels. For each voxel set, we conducted a 2 (Script: English, Chinese) × 3 (Group: L2 pre-readers, L2 beginning readers, L2 advanced readers) mixed-design ANOVA, with Script as a within-subject factor and Group as a between-subject factor. The Script × Group interaction was used to assess whether the relative response profile for English and Chinese words differed across stages of L2 literacy acquisition. To further characterize stage-related differences in responses to each script, one-way ANOVAs were conducted for English and Chinese beta estimates across groups. Linear trend analyses were also performed using GroupCode, coded as 1, 2, and 3 for L2 pre-readers, beginning readers, and advanced readers, respectively.

Under the Overlay Model, English responses were expected to increase with L2 literacy experience, while Chinese responses were expected to remain stable. In contrast, the Competing Model predicted that increasing English responses would be accompanied by reduced Chinese responses, reflecting competitive reallocation within the VWFA.

#### 2.5.4 Representational similarity analysis (RSA) in the VWFA

Building on the preceding univariate activation analyses, we next examined how L2 literacy acquisition shapes the representational structure of L1 and L2 words in the VWFA. Specifically, we conducted RSA to assess multivariate patterns of L1 and L2 word representations within the VWFA in each participant, and to compare these patterns across the three groups.

Given the mini-block design and the inclusion of a brief attention task (star detection) within each block, RSA was conducted on unsmoothed functional data, with block-level activation estimates computed by excluding star-related time periods to isolate stimulus-driven responses. Within a 10-mm-radius sphere centered on the classic VWFA coordinates, β estimates for each block were extracted and used to construct a 16×16 representational similarity matrix (4 conditions×4 blocks). Each cell of the matrix reflected the Pearson correlation between activation patterns across blocks. To reduce potential inflation of similarity estimates due to run-specific noise or temporal autocorrelation, only cross-run correlations were included in the analysis. The same RSA procedure was repeated using 6- and 8-mm-radius spheres centered on the same VWFA coordinates to assess the robustness of the findings.

To characterize how L2 literacy experience shapes L2 word representations and their relationship to L1 word representations in the VWFA (overlay vs. competing), we examined three sets of comparisons. All correlation coefficients were Fisher z-transformed prior to analysis. First, we assessed whether the VWFA exhibited robust representations of each script at each stage of L2 reading acquisition. A robust word representation was operationally defined as greater within-category similarity (English–English or Chinese–Chinese) than the mean similarity between that script and the non-linguistic categories (English–faces and English–houses, or Chinese–faces and Chinese–houses) within each group. Second, we examined how within-script similarity for English (English–English) and Chinese (Chinese–Chinese) each differed across groups using separate one-way ANOVAs, to assess whether L2 word representations became increasingly robust while L1 word representations remained stable or declined. Third, we examined the representational relationship between L1 and L2 words by comparing within-script similarities (English–English, Chinese–Chinese) with cross-script similarity (English–Chinese). Within each group, paired-sample t-tests were used to assess whether L1 and L2 converged toward a shared representational structure or remained distinct at each reading stage. For comparisons of within-script versus script-non-linguistic similarity, FDR correction was applied across the three group-wise paired tests for each script. For comparisons among English–English, Chinese–Chinese, and English–Chinese similarity, FDR correction was applied across the three pairwise tests within each group.

#### 2.5.5 Supervised machine learning classification based on activation patterns in the VWFA

Additionally, we applied supervised machine learning to assess whether VWFA activation patterns evoked by different stimulus categories, particularly English words and Chinese words, could reliably differentiate participants with varying L2 literacy levels (L2 pre-readers, beginning readers, and advanced readers). For each participant, GLM-derived β estimates for each stimulus category (English words, Chinese words, faces, and houses) relative to fixation were extracted from a 6-mm spherical ROI centered on the canonical VWFA coordinates. This relatively small ROI was selected to restrict the number of voxel features and thereby reduce the likelihood of overfitting. Voxel-wise β values from this ROI were obtained for each run, yielding a feature matrix of size (subjects ×runs) × voxels for each stimulus category. Each sample was labeled according to the participant’s group (L2 pre-readers, beginning readers, or advanced readers) (**Fig. 5A**).

**Fig. 4.**
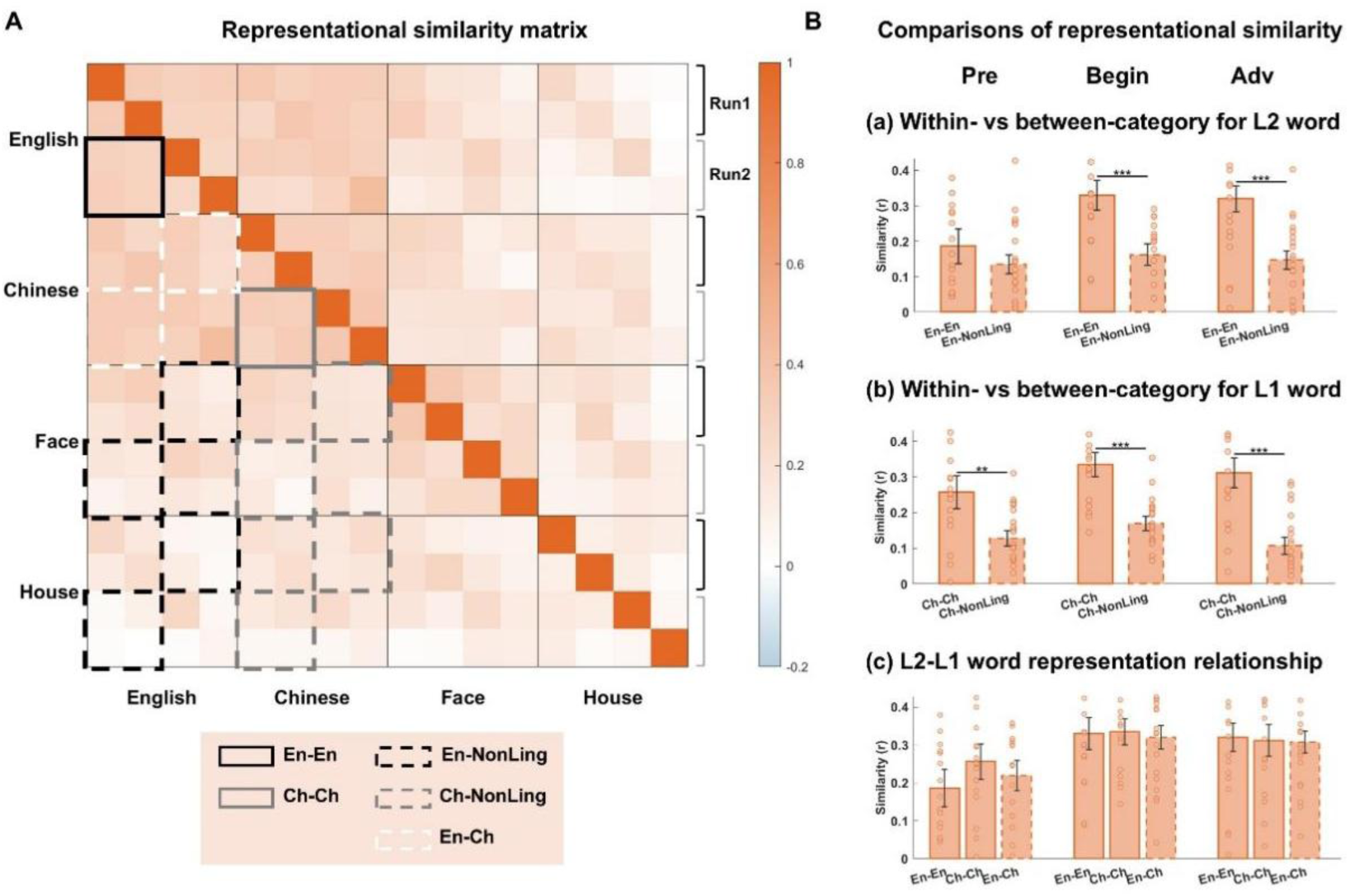
Representational similarity analyses of word representations in the VWFA. (A) Representational similarity matrix (RSM) showing the mean, across participants, of the pairwise correlations between spatial activation patterns evoked by four stimulus categories: English words, Chinese words, faces, and houses. Color intensity indicates the magnitude of similarity (warmer colors = higher similarity; cooler colors = lower similarity). Solid boxes indicate within-category similarities (En-En, Ch-Ch), and dashed boxes indicate between-category similarities (En-NonLing, Ch-NonLing, En-Ch). En-NonLing (Ch-NonLing) denotes the mean between-category similarity between English (Chinese) words and non-linguistic stimuli (faces and houses). (B) Comparisons of representational similarity in the VWFA. Bar plots show mean similarity values for three comparisons: (top) within-English similarity versus English-non-linguistic similarity, (middle) within-Chinese similarity versus Chinese-non-linguistic similarity, and (bottom) comparisons among within-English, within-Chinese, and English-Chinese similarity. Results are shown separately for the three groups differing in L2 experience (pre-, beginning, and advanced readers). Significance levels: \**p_FDR_* < .05, \*\**p_FDR_* < .01, \*\*\**p_FDR_* < .001 (FDR-corrected paired-sample t-tests).

**Fig. 5.**
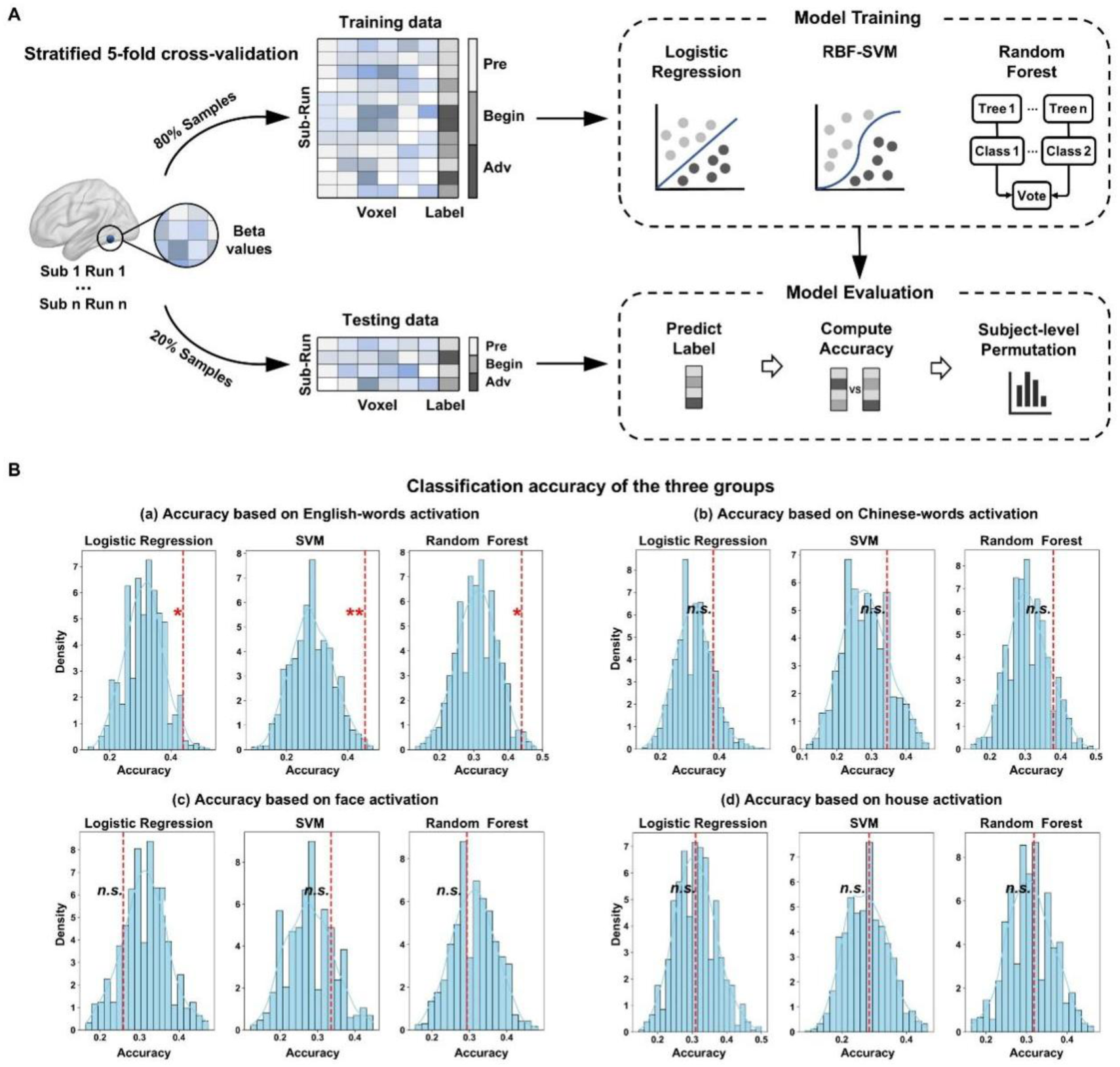
Classification of L2 literacy stage based on VWFA activation patterns. (A) Schematic of the supervised machine learning pipeline. For each participant, GLM-derived β estimates for each stimulus category were extracted from a 6-mm spherical ROI centered on canonical VWFA coordinates. Voxelwise β values from each run formed a feature matrix of size (subjects×runs)×voxels, with each sample labeled according to L2 proficiency group (pre-readers, beginning readers, advanced readers). Three classifiers (logistic regression, support vector machine with radial basis function kernel, and random forest) were evaluated using a five-fold StratifiedGroupKFold cross-validation procedure, which ensured that (i) samples from the same participant were not split across training and test sets, and (ii) class proportions were approximately balanced across folds. Classification accuracy was computed by aggregating predictions across all folds. Statistical significance was assessed using permutation testing with participant-level label shuffling. The full cross-validation procedure was repeated 1,000 times to generate a null distribution of classification accuracies. (B) Classification performance based on VWFA activation patterns for each stimulus category. Blue histograms represent the null distribution of classification accuracies from 1,000 permutations of group labels. The vertical red line indicates the observed classification accuracy. Significance levels: \**p* < .05, \*\**p* < .01; n.s., not significant.

We implemented a stratified group-based cross-validation procedure. Specifically, five-fold StratifiedGroupKFold cross-validation was used to ensure that (i) samples from the same participant were always assigned to the same fold and never split between the training and test sets, and (ii) class proportions were approximately balanced across folds. Within each fold, feature standardization was performed using a StandardScaler fitted to the training data and subsequently applied to the test data. Three classifiers were trained on the training set and evaluated on the held-out test set: logistic regression, support vector machine (SVM with an RBF kernel), and random forest. Classification accuracy was computed by pooling predictions for all held-out samples across the five folds. Statistical significance was assessed using permutation testing with participant-level label shuffling. In each permutation, group labels were shuffled across participants, and the same shuffled label was assigned to both runs from a given participant. This preserved the repeated-measures structure of the data. The full five-fold cross-validation procedure was then repeated using the shuffled labels. A total of 1,000 permutations were performed to generate a null distribution of classification accuracies. The permutation-based *p* value was calculated as (*k* + 1)/(*N_perm_* + 1), where *k* was the number of permuted accuracies equal to or greater than the observed accuracy.

#### 2.5.6 Analyses of other spoken-language-related regions

Given that reading experience is known to reshape not only the ventral visual pathway but also broader components of the distributed language network (Dehaene et al., 2010; Monzalvo & Dehaene-Lambertz, 2013), we conducted additional analyses in several left-hemisphere spoken-language-related regions. Guided by an integrative review and prior meta-analyses of reading-related activation (Houdé et al., 2010; A. Martin et al., 2015; Romanovska & Bonte, 2021; Walenski et al., 2019), we defined four 6-mm-radius spherical ROIs across the left frontal, parietal, and temporal cortices: the middle frontal gyrus (MFG), intraparietal sulcus (IPS), superior temporal gyrus (STG), and superior temporal sulcus (STS) **(Fig. 6A)**.

**Fig. 6.**
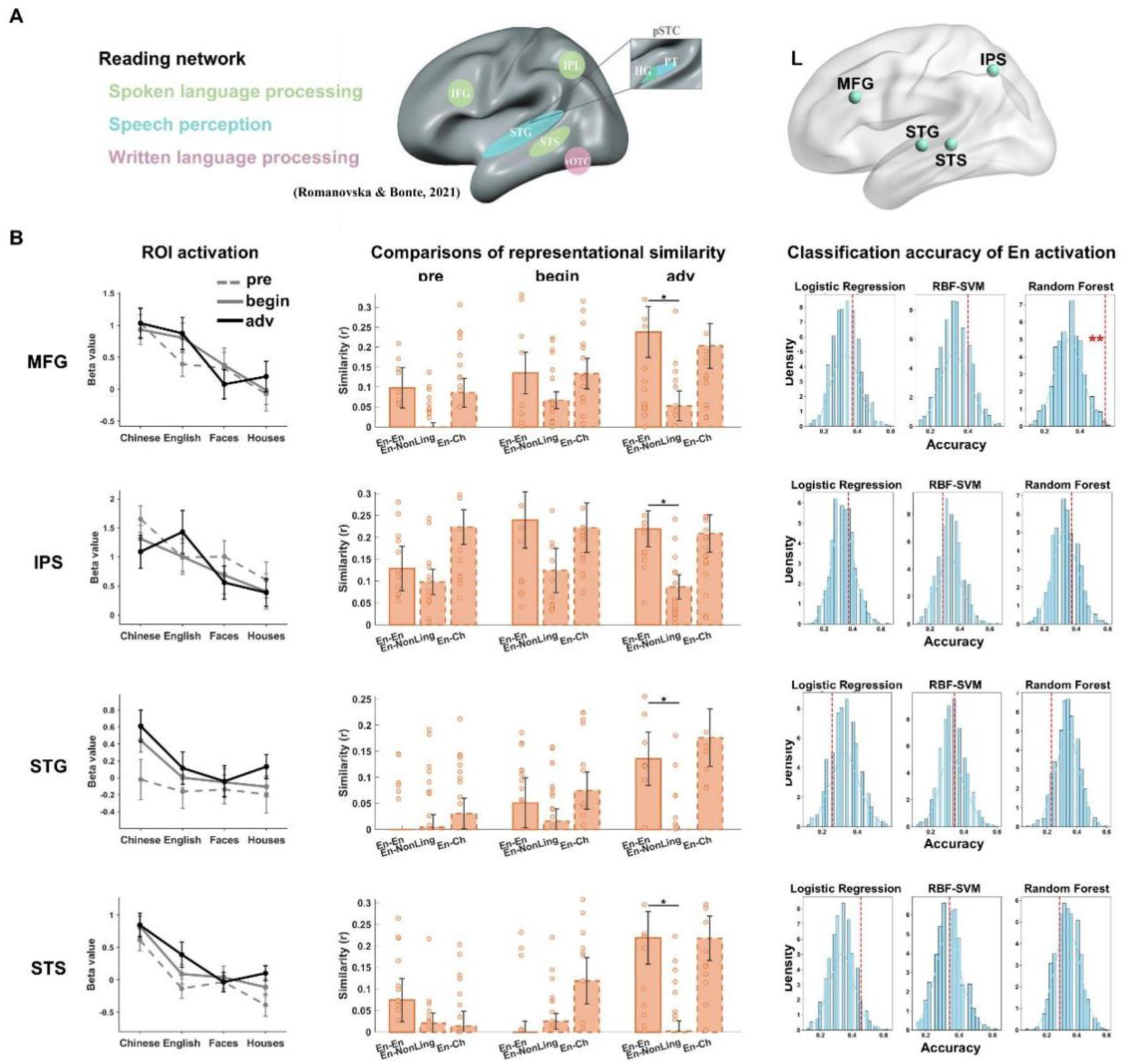
ROI activation, RSA, and supervised machine learning classification analyses of left spoken-language-related regions. (A) Illustration of the four spoken-language-related ROIs in the left frontal, parietal, and temporal cortex, including the left MFG, IPS, STG, and STS, defined based on prior review and meta-analyses of reading. (B) Results are organized into three panels, with each row corresponding to one ROI and each column representing a distinct analysis. **Left panel**: Univariate activation for four categories (English words, Chinese words, faces, houses) across the three groups. **Middle panel**: Comparisons of representational similarity separately for three groups. Bar plots show mean similarity values for two comparisons: within-English similarity versus English-non-linguistic similarity and English-Chinese similarity. **Right panel**: Classification performance for three classifiers trained to predict L2 group membership from English-word activation patterns. Blue histograms represent the null distribution of classification accuracies from 1,000 permutations with participant-level label shuffling. The vertical red line indicates the observed accuracy aggregated across cross-validation folds. Significance levels: \**p* < .05, \*\**p* < .01, \*\*\**p* < .001.

First, we performed univariate analyses within each ROI to examine activation magnitudes for each stimulus category across stages of L2 literacy acquisition. For each participant, GLM-derived β estimates were extracted for the four visual categories (English words, Chinese words, faces, houses) versus fixation. One-way ANOVAs were conducted for each stimulus category to examine group differences among pre-, beginning, and advanced readers.

Second, we performed RSA within each spoken-language-related ROI following the same procedure used for the VWFA (Section 2.5.4). Block-level β estimates were extracted from each ROI, and cross-run representational similarity matrices were constructed for each participant. Focusing on L2 word representations in the broader language network, we compared z-transformed within-English similarity with (1) the similarity between English words and non-linguistic categories and (2) the similarity between English and Chinese words within each ROI using paired-sample t-tests. FDR correction was applied separately within each ROI across the six planned comparisons, corresponding to two comparisons in each of the three reading groups. These analyses tested the robustness of English-word representations and their relationship to corresponding Chinese-word representations.

Finally, we applied supervised machine learning classification to evaluate whether English-word activation patterns in these spoken-language-related regions could differentiate participants with varying L2 reading experience. Following the procedure used for the VWFA (Section 2.5.5), GLM-derived run-level β estimates for English words were extracted from each ROI and used as input features. The classifiers were trained and evaluated separately for each ROI using five-fold StratifiedGroupKFold cross-validation. Classification accuracy was calculated by pooling predictions for all held-out samples across the five folds. Statistical significance was assessed using permutation testing with participant-level label shuffling.

## 3 Results

### 3.1 Category-selective activation at the whole-brain level

To characterize category-selective activation profiles across the ventral visual cortex, we first examined responses across all children. The four categories showed a broadly medial-to-lateral organization within the vOTC: house-selective activation was bilateral and centered in medial occipitotemporal cortex, spanning calcarine and lingual visual cortices as well as parahippocampal cortex; face-selective activation was predominantly right-lateralized, occupying adjacent fusiform and occipitotemporal regions; and Chinese- and English-word-selective activations were primarily left-lateralized (**Fig. 1A**). Beyond the left ventral activations, Chinese-word-selective activation extended to the left frontal regions (precentral gyrus, middle frontal gyrus and supplementary motor area), left superior and middle temporal regions, as well as right superior temporal gyrus. English-word-selective activation extended to the left frontal regions (precentral gyrus, superior and inferior frontal gyrus), together with the inferior parietal lobule and inferior temporal gyrus (**Supplementary Table S2**). For non-linguistic categories, face-selective activation extended into bilateral fusiform cortex, with additional right inferior occipital and lingual activation and medial temporal clusters involving the parahippocampal regions. House-selective activation was extensive and bilateral across the fusiform gyrus, middle occipital gyrus, calcarine cortex, and parahippocampal cortex.

We next examined activation patterns within each group (**Fig. 1B–D**). As predicted, a preferential response to Chinese words in the left vOTC was consistently observed across all three groups, indicating stable L1-word selectivity regardless of L2 experience. In contrast, activation to English words in the left vOTC showed different profiles across groups: pre-readers showed limited English-selective activation, beginning readers showed English-selective voxels emerging in the vicinity of Chinese-selective regions, and advanced readers showed substantial spatial overlap between English- and Chinese-selective voxels. Activation patterns for faces and houses remained largely consistent across the three groups.

We further conducted a one-way ANOVA and a whole-brain multiple regression analysis to investigate group differences and the association between L2 literacy proficiency and L2-related activation, respectively. We first tested for these effects using a stringent whole-brain threshold (*p* < .001 at the voxel level and *p* < .05 FWE-corrected at the cluster level), which yielded no significant clusters. For exploratory visualization and hypothesis generation, we further inspected the data at a less stringent threshold (*p* < .001 and *p* < .005 at the voxel level, cluster-level uncorrected). This analysis identified substantially overlapping voxels in the left posterior occipitotemporal cortex, showing both a main effect of group in the ANOVA and an association with the English literacy composite score in the regression analysis (**Supplementary Fig. S1**). Although these uncorrected findings should be interpreted cautiously, their spatial distribution was consistent with our prior focus on the vOTC encompassing the VWFA.

### 3.2 Moving-window ROI activations along the lateral–medial and posterior–anterior axes centered on the VWFA

To examine category-selective activation along the lateral–medial axis, with a focus on responses to English and Chinese words, we kept y = –57 and z = –12 constant and extracted activation for English words, Chinese words, faces and houses along the x-axis (ranging from –57 to –21; **Fig. 2A**). Along the lateral–medial axis, the activation profiles showed that Chinese words elicited consistent peak activation at the canonical VWFA coordinate (Vogel et al., 2012), x = –45, across all three groups, indicating robust L1-word responses regardless of L2 experience. In contrast, English-word responses at this coordinate were weak in L2 pre-readers but were comparable to Chinese-word responses in beginning and advanced readers (**Fig. 2B**). In L2 pre-readers, the English–Chinese difference at x = –45 was nominally significant before correction but did not survive FDR correction (*t* = 2.67, *p*_uncor_ = .015, *p*_FDR_ = .182). As L2 experience accumulated, both beginning and advanced readers exhibited comparable peak responses to English and Chinese words at x = –45, suggesting that L2-word responses emerged within the same general VWFA territory and exhibited a spatial profile aligned with that of L1 responses, a pattern consistent with the Overlay Model. The non-linguistic categories showed the expected topographic pattern, with faces and houses peaking at relatively medial coordinates across all three groups.

Consistent with these descriptive patterns, word-selectivity analyses along the lateral–medial axis further corroborated these findings: one-way ANOVAs revealed a significant group effect for English-word selectivity at x = –45 (*F*(2, 55) = 3.28, *p* < .05, η² = .11). Post hoc comparisons indicated that L2 pre-readers showed lower English-word selectivity than both beginning and advanced readers (**Fig. 2C**). No group effect was observed for Chinese-word selectivity at any coordinate along this axis, and no other lateral–medial ROI showed group differences in English-word selectivity.

Along the posterior–anterior axis (x = –45, z = –12; y = –72 to –24; **Fig. 2D**), the activation profiles showed that L2 pre-readers exhibited nominally lower English than Chinese activation at y = –72 (*t* = 2.88, *p*_uncor_ = .010, *p*_FDR_ = .144) and y = –60 (*t* = 2.65, *p*_uncor_ = .016, *p*_FDR_ = .118), coordinates slightly posterior to the classic VWFA location (**Fig. 2E**). In contrast, beginning and advanced readers showed comparable responses to English and Chinese words throughout this axis.

Complementing these gradient plots, word-selectivity analyses along the posterior–anterior axis showed a nominal group effect for English-word selectivity at y = –72 (*F*(2, 55) = 3.39, *p* < .05, η² = .11). Post hoc comparisons showed that L2 pre-readers exhibited lower English-word selectivity than both beginning and advanced readers (**Fig. 2F**). Chinese-word selectivity showed a nominal group effect at y = –36 (*F*(2, 55) = 3.92, *p* < .05, η² = .12), with higher Chinese-word selectivity in L2 pre-readers than in advanced readers. No other ROIs along the posterior-anterior axis exhibited significant group differences in either English- or Chinese-word selectivity.

In summary, these results delineate the spatial reorganization associated with L2 literacy acquisition along the lateral–medial and posterior–anterior axes within and around the VWFA. Notably, in L2 beginning and advanced readers, activation profiles for Chinese and English words followed closely aligned spatial gradients along both axes, with peak responses occurring at the same sampled coordinates and reaching comparable magnitudes. By contrast, L2 pre-readers showed a broadly similar spatial profile, but English-word responses were weaker than Chinese-word responses at the shared peak locations, resulting in less pronounced English-word selectivity. These findings suggest that L2 reading recruits the same general VWFA territory as L1 processing. Importantly, this regional alignment was accompanied by stable L1-evoked responses across groups, consistent with the Overlay Model.

### 3.3 Individual-level analyses: Cross-run responses in the most responsive VWFA voxels

To complement the group-level and ROI-based analyses, which assume consistent functional localization across children, we examined cross-language responses within the VWFA at the individual level. Using a cross-run selection-extraction procedure, we identified for each child the VWFA voxels most responsive to English and Chinese words and extracted independent beta estimates for both scripts from these voxel sets. This analysis tested whether L2 responses increased with L2 literacy experience within individually defined top-responsive VWFA voxels, and whether this increase was accompanied by any reduction in native-script responses.

In English top-10% voxels, English responses increased across groups, whereas Chinese responses remained relatively stable (**Fig. 3B**). The 2 × 3 mixed-design ANOVA revealed a significant Script × Group interaction, *F*(2, 55) = 3.39, *p* = .041, η_p_² = .11. Within-group paired comparisons showed that only L2 pre-readers showed a significant script difference, with lower English than Chinese responses. Script-specific one-way ANOVAs showed a significant group effect for English beta, *F*(2, 55) = 4.68, *p* = .013, η² = .15, with lower English responses in L2 pre-readers than in the two reader groups. Consistent with this pattern, the linear trend analysis showed a significant positive linear trend in English beta estimates across the ordered L2 literacy groups, *b* = 0.30, *t*(56) = 2.62, *p* = .011. In contrast, Chinese beta did not show a significant group effect, nor a significant linear trend.

In Chinese top-10% voxels, the 2 × 3 mixed-design ANOVA did not reveal a significant Script × Group interaction. Script-specific analyses showed that English beta did not differ significantly across groups, although the linear trend analysis revealed a significant positive linear trend across the ordered L2 literacy groups, *b* = 0.23, *t*(56) = 2.32, *p* = .024. Chinese beta showed no significant group effect, and no significant linear trend.

Together, these individual-level analyses provide converging evidence that L2 literacy acquisition was not accompanied by a reduction in L1 responses within the VWFA. The clearest group-related difference was observed in English top-10% voxels, where English responses increased from pre-readers to readers while Chinese responses showed no evidence of group differences. In Chinese top-10% voxels, L1 responses likewise showed no significant group difference, and there was no significant Script × Group interaction. This pattern supports the Overlay Model, suggesting that the VWFA incorporates a newly learned script primarily through enhanced L2 responses in English-responsive voxels, without detectable attenuation of native-script responses.

### 3.4 Representational similarity analysis (RSA)

To examine whether learning to read L2 shapes the internal representational structure of the VWFA beyond differences in activation magnitude, we conducted RSA within this region. For each participant, a representational similarity matrix was computed based on the cross-run correlations between voxel-wise activation patterns evoked by the four stimulus categories (English words, Chinese words, faces, and houses) within the VWFA sphere (**Fig. 4A**). We compared within-category similarity (e.g., En–En) with the mean between-category similarity (e.g., En–faces/houses) to assess whether the VWFA exhibited robust representations of L1 and L2 words at each reading stage (**Fig. 4B, panels a–b**), examined how within-category similarity for English and Chinese words each differed across groups, and compared within-script with cross-script similarities to characterize the representational relationship between L1 and L2 (**Fig. 4B, panel c**).

Comparisons between within-category similarity and the mean between-category similarity for English words (En-En vs En-faces, En-houses) revealed that beginning and advanced readers exhibited significantly higher within-category similarity (beginning readers: *t* = 3.96, *p*_FDR_ < .001, Cohen’s *d* = 0.91; advanced readers: *t* = 4.18, *p*_FDR_ < .001, Cohen’s *d* = 0.96), whereas pre-readers did not show a reliable difference (**Fig. 4B, panel a**). In contrast, comparisons between within-category similarity and the mean between-category similarity for Chinese words (Ch-Ch vs Ch-faces, Ch-houses) showed consistently higher within-Chinese similarity across all three groups (pre-readers: *t* = 3.24, *p*_FDR_ < .01, Cohen’s *d* = 0.73; beginning readers: *t* = 6.13, *p*_FDR_ < .001, Cohen’s *d* = 1.41; advanced readers: *t* = 5.38, *p*_FDR_ < .001, Cohen’s *d* = 1.23) (**Fig. 4B, panel b**). For the comparison among within-English, within-Chinese, and English-Chinese similarity, FDR-corrected pairwise paired-sample t-tests revealed no significant differences in any group (**Fig. 4B, panel c**). One-way ANOVAs further showed no significant group effect for within-English similarity (*F*(2, 55) = 2.78, *p* = .071, η² = .09), or within-Chinese similarity (*F*(2, 55) = 0.91, *p* = .411, η² = .03). Analyses using 6-mm and 8-mm-radius spheres centered on the canonical VWFA coordinates replicated these findings (**Supplementary Fig. S2**).

Taken together, these results suggest that robust L2 word representations emerged only in children with L2 reading experience, while robust L1 word representations were evident across all three groups. Within-category similarity for Chinese words did not show a significant group effect, with no evidence of L1 word representational decline as predicted by the Competing Model. Although within-category similarity for English words did not differ significantly across groups, reliable L2 word representations were observed in L2 beginning and advanced readers, but not in L2 pre-readers, indicating an experience-dependent emergence of L2 word representations within the VWFA.

### 3.5 Supervised machine learning classification

To assess whether VWFA responses to different stimulus categories could differentiate participants with varying levels of L2 reading experience, we conducted supervised machine learning analyses based on run-level GLM-derived β estimates. Results revealed that VWFA activation patterns to English words reliably discriminated participants across the three groups, with classification accuracies significantly above chance for all three models: logistic regression (accuracy = .44, *p* = .024), support vector machine (accuracy = .46, *p* = .008), and random forest (accuracy = .44, *p* = .019). In contrast, classification based on VWFA responses to Chinese words, faces, or houses did not exceed chance level in any classifier (*ps* > .10). These findings indicate that VWFA responses to L2 words change systematically with reading acquisition, enabling reliable group discrimination, whereas responses to L1 words and non-linguistic categories did not contain sufficient information for above-chance group classification. This pattern is consistent with the Overlay Model, suggesting that VWFA reorganization is driven by emerging L2 word representations rather than reallocation of L1 resources.

### 3.6 Analyses of other language-related regions

To characterize how L2 reading experience shapes the functional organization of the left spoken-language network, we investigated these language-related ROIs (STS, STG, IPS, MFG, see **Fig. 6A**) using the analytical framework established for the VWFA, which combines univariate activation, RSA, and supervised classification.

Univariate analyses revealed no significant group effects for any stimulus category in any of the four ROIs (*ps* > .05), indicating no reliable differences in mean activation magnitude among L2 pre-readers, beginning readers, and advanced readers. Descriptive patterns across the four ROIs suggested region-specific differences in English-word responses associated with L2 reading experience (**Fig. 6B, left panel**). RSA analyses revealed that no robust L2 word representations were observed in pre-readers or beginning readers after FDR correction, as their within-English similarity did not exceed English-non-linguistic similarity in any ROI. In advanced readers, by contrast, within-English similarity exceeded English-non-linguistic similarity in the MFG (*t* = 3.48, *p*_FDR_ < .05, Cohen’s *d* = 0.80), STG (*t* = 3.38, *p*_FDR_ < .05, Cohen’s *d* = 0.77), STS (*t* = 3.38, *p*_FDR_ < .05, Cohen’s *d* = 0.77), and IPS (*t* = 4.41, *p*_FDR_ < .05, Cohen’s *d* = 1.01), indicating that robust L2 word representations in language-related ROIs were detectable only in advanced readers (**Fig. 6B, middle panel**). In addition, no ROI showed significantly greater within-English than English-Chinese similarity after FDR correction. Finally, supervised classification analyses demonstrated that random-forest classification based on MFG patterns exceeded chance (accuracy = .54, *p* = .008), whereas classification using logistic regression or an SVM did not. More broadly, none of the language-related ROIs could consistently decode L2 reading stages across all three classifiers (**Fig. 6B, right panel**). These decoding results suggest that, compared to the visual cortex, these spoken-language regions generally lack sufficient group-discriminative information to be sensitive to variations in L2 reading experience.

## 4. Discussion

The present study investigated how the VWFA reorganizes to incorporate a structurally distinct second writing system during early L2 literacy acquisition. Using a cross-sectional design, native Mandarin-speaking children were divided into three groups by L2 experience: L2 pre-readers, beginning readers, and advanced readers. Whereas pre-readers showed little evidence of robust L2-word selectivity, beginning readers exhibited reliable L2-word selectivity within the same general VWFA territory already responsive to L1 words, with this pattern remaining stable in advanced readers. Representational similarity analyses further revealed robust L2 word representations in beginning and advanced readers, but not in pre-readers, indicating that L2 literacy acquisition drives the integration of L2 word representations into a VWFA region already committed to L1 word processing. To characterize the nature of this reorganization, we first examined how L2-selective responses, defined as greater activation to L2 relative to other non-linguistic categories, arose in the VWFA. We found that the selective responses were driven by increased activation to L2 stimuli rather than by decreased activation to L1 or other non-linguistic categories. We then assessed whether the emergence of L2 word representations disrupted pre-existing L1 word representations and found no evidence of disruption, indicating that L2 acquisition did not degrade prior L1 responses. Converging evidence from machine learning analyses further revealed that only L2 responses, but not L1 or other category responses, reliably discriminated among the three L2 literacy groups. Taken together, these findings indicate that early L2 literacy acquisition prompts the incorporation of representations of a newly learned script into an already specialized VWFA without measurable competition with either pre-existing L1 word representations or non-linguistic category representations (e.g., faces, houses), thus supporting an Overlay Model.

### 4.1 Neural reorganization induced by learning to read

The present study extends prior work on native-script acquisition to the domain of L2 reading, revealing similarly reorganization of the VWFA in response to a newly learned script. This parallels evidence from native-script learning demonstrating that even a few weeks to months of reading instruction can induce measurable changes in the ventral occipitotemporal cortex (Brem et al., 2010; Dehaene-Lambertz et al., 2018; Yeatman et al., 2024). Consistent with this, our previous study comparing 6-year-old L1 pre-readers, 6-year-old beginning readers, and 9-year-old advanced readers demonstrated that such reorganization is driven by reading experience rather than maturation (Feng et al., 2022). While a growing body of research has examined the neural correlates of L2 reading acquisition, particularly how the VWFA supports the learning of a second writing system, these studies have focused predominantly on adults learning artificial orthographies (Martin et al., 2019; Moore et al., 2014; Xue et al., 2006), leaving it largely unknown how the developing brain incorporates L2 word representations into the existing reading network. The present findings directly address this gap, showing that in children with well-established L1 word representations in the VWFA, this region remains sufficiently plastic to support experience-dependent reorganization for L2 literacy acquisition. Together, these findings suggest that acquiring a native script does not exhaust the brain’s plasticity; rather, the VWFA retains the capacity for continued experience-dependent reorganization in response to additional writing systems during development.

### 4.2 Absence of neural competition between L1 and newly acquired L2 in the VWFA

A key prediction of the Competing Model, informed by the cortical recycling hypothesis, is that learning to read a new script requires the reallocation of cortical resources, potentially encroaching upon territory previously dedicated to pre-existing visual categories (Dehaene & Cohen, 2007). In the context of L2 literacy acquisition, this view predicts that the emergence of L2 responses would trigger neural competition, degrading or displacing pre-existing L1 responses within the shared cortical territory of the VWFA. Our findings, however, provide no evidence in support of this competitive encroachment account. Instead, the VWFA appears to incorporate L2 print processing without measurably disrupting pre-existing L1 responses or representations, suggesting that intra-category coexistence, rather than competition, characterizes the early stages of L2 literacy acquisition.

First, the incorporation of L2 script into the VWFA did not occur at the expense of native-script processing. Both univariate activation and RSA revealed that L1-word responses remained remarkably stable across all three groups. Specifically, the magnitude of L1-selective activation did not decrease from L2 pre-readers to advanced readers, and intra-category pattern similarity for L1 words showed no evidence of degradation. Crucially, classification based on L1 activation patterns did not reliably distinguish among the three groups. This clear preservation of L1 sensitivity directly argues against the notion of neural competition between the two writing systems within the VWFA, indicating that the existing neural substrate possesses sufficient capacity to incorporate an additional, structurally distinct script through an overlay mechanism rather than a competing one.

These findings stand in contrast to prior evidence implicating competitive dynamics in reading-related cortical organization. In studies of illiterate adults, higher reading ability has been associated with a modest reduction in face responsivity within the left fusiform gyrus (Dehaene et al., 2010). Similarly, struggling young readers exhibit lower word selectivity in the VWFA not due to diminished responses to print, but due to a failure to suppress competing responses to non-linguistic objects (Kubota et al., 2019). More recently, longitudinal evidence has demonstrated that increases in word selectivity during childhood are directly coupled with decreases in limb selectivity within the ventral temporal cortex, suggesting that limb-selective cortex is progressively repurposed for orthographic processing (Nordt et al., 2021). Going beyond these cross-category competitions, our study investigated whether a parallel competitive dynamic exists when acquiring a new script. We found no such displacement within the VWFA. The robust increase in L2-word activation observed in beginning and advanced readers was accompanied by stable responses to Chinese words, and activation profiles for non-linguistic categories within the VWFA remained stable across groups. This absence of competition may partly reflect the fact that writing systems, despite their surface diversity, share a common set of basic visual features, such as lines and angles at particular junctions, that were culturally selected to match contours recurring in natural scenes (Changizi et al., 2006). This shared low-level structure may allow the neural circuitry recruited for L1 print processing to accommodate a second, visually distinct script with minimal need for additional resources.

Taken together, our cross-sectional findings suggest that when an additional script is introduced after the L1 reading network is already established, it seamlessly overlays onto the existing word-selective architecture of the VWFA without triggering cortical encroachment upon existing L1 representations. Although the absence of limb stimuli in the current paradigm precludes a direct voxel-level test of the limb-displacement hypothesis (see Limitations), the overall stability of both L1 and non-linguistic category representations within the VWFA strongly implies that L2 print exploits pre-existing orthographic channels carved out during L1 acquisition, reducing the need for further competitive reallocation of the broader visual cortex.

### 4.3 L2 literacy acquisition recruits the existing VWFA territory: accommodation and assimilation

In addition to asking whether the VWFA would undergo systematic modifications as children begin acquiring a second script, we further examined the nature of this cortical reorganization. We first examined how this reorganization was spatially structured within the VWFA. A key prediction of the Overlay Model is that L2 print may recruit the same cortical territory already tuned to the native script. Consistent with this prediction, spatial activation patterns showed that Chinese words consistently evoked stronger responses than non-linguistic categories within the left ventral visual cortex across all three groups, indicating that selective cortical responses to the native script were already well established. Fine-grained moving-window ROI analyses further revealed that English-word responses near the VWFA were weak in L2 pre-readers, but grew and peaked within the same general territory once children acquired English reading experience, giving rise to L2 responses that closely aligned with L1 responses in both magnitude and spatial location. Thus, the critical spatial change associated with L2 literacy acquisition was the emergence of L2-word selectivity within pre-existing VWFA territory despite already robust native-script specialization. Although this regional alignment does not imply exact voxel-by-voxel identity, the overall pattern supports the view that L2 is spatially incorporated within a shared cortical territory.

This spatial pattern is consistent with adult bilingual studies showing that two writing systems can engage overlapping occipitotemporal regions, including studies of English–Hebrew, Chinese–Korean, and Chinese–English bilingual readers (Bai et al., 2011; Baker et al., 2007; Wong et al., 2009). The present study extends these adult findings by showing that, even from the earliest stages of L2 literacy acquisition, a structurally distinct writing system is incorporated into the existing VWFA territory rather than assigned to a separate cortical substrate. This interpretation is further supported by evidence from training studies, where adults learning artificial logographic systems likewise recruit neural territory previously specialized for native-script literacy (Martin et al., 2019).

Moreover, in the literature on bilingual reading, assimilation and accommodation have been proposed to account for how the brain adapts to L2 learning (Perfetti et al., 2007). When L2 is similar to L1, the brain recruits the same network already established for L1 reading, a process termed assimilation, as demonstrated in fMRI studies of English–Korean bilinguals, in which both languages use alphabetic writing systems (Kim et al., 2016). When L2 is substantially different from L1, the brain recruits additional or distinct regions to support L2-specific processing demands, a process termed accommodation, as suggested in studies of English L1 participants learning Chinese as L2, including recruitment of the right fusiform region, middle occipital gyri, and middle frontal gyrus (Liu et al., 2007; Nelson et al., 2009). The substantial structural differences between Chinese and English might lead one to predict accommodation in Chinese–English bilingual readers. Our results, however, reveal that the VWFA specifically follows an assimilation pattern, with comparable responses to both L1 and L2 observed in beginning and advanced readers. We argue that this reflects the core functional role of the VWFA: although Chinese and English differ substantially at many levels of linguistic representation, both writing systems share the fundamental demand of orthographic processing. Prior research has consistently shown that the VWFA is sensitive to orthographic structure in both languages, as evidenced by a shared lexicality gradient whereby responses systematically increase as stimuli more closely approximate real words (Zhan et al., 2023). Since this orthographic processing demand is common to both scripts, the VWFA is well positioned to assimilate L2 within its existing L1 framework, regardless of the broader structural differences between the two writing systems.

### 4.4 Beyond the VWFA: gradual integration of L2 into the broader spoken-language network

In the present study, we applied the same analytical framework used for the VWFA—incorporating univariate activation, RSA, and supervised classification—to examine how regions within the left spoken-language network (including the MFG, IPS, STG, and STS) were recruited during L2 reading acquisition. In contrast to the reorganization observed in the VWFA, these spoken-language regions exhibited a less immediate and more heterogeneous L2-sensitive profile. Descriptively, across nearly all groups, these regions showed a general tuning for written language, responding more strongly to words than to non-linguistic stimuli. English words also tended to elicit weaker activation than Chinese words across most groups and ROIs, suggesting that the spoken-language network remained more strongly engaged by the native script. Moreover, whereas robust L2 word representations emerged in the VWFA in beginning readers, comparable representations in these speech-related regions arose predominantly in advanced readers, suggesting a prolonged trajectory (**Fig. 6B**). Aligning with this more gradual pattern, supervised classification revealed that spoken-language regions generally lacked sufficient group-discriminative information to reliably decode L2 reading stages.

This asymmetry likely reflects the sequential process through which a newly learned writing system becomes integrated into the reading network. The VWFA is directly engaged by visual print exposure and orthographic learning, and may therefore develop sensitivity to L2 literacy experience relatively early (Brem et al., 2010; Dehaene-Lambertz et al., 2018). By contrast, spoken-language-related regions may rely on the gradual coupling of orthographic representations with phonological, articulatory, and lexical-semantic processes, which likely require more extended reading experience to consolidate (Dehaene-Lambertz et al., 2018). The present findings therefore extend prior evidence that reading acquisition reshapes both visual and spoken-language systems, now in the context of L2 literacy, and further suggest that these two components do not reorganize in tandem. Specifically, L2 print appears to be initially incorporated within the VWFA, whereas the functional engagement of broader language circuits unfolds more gradually.

Crucially, the protracted nature of this neural integration may be profoundly shaped by the characteristics of the L2 writing system. This notion is consistent with previous cross-linguistic evidence showing that the degree of orthographic transparency modulates the timing and pace of this integration. More specifically, in a transparent language like Polish, print–speech convergence was observed at an early stage of reading acquisition (Dębska et al., 2024), while in an opaque language like English, the convergence was observed in a more gradual fashion (Chyl et al., 2021). Although these findings were originally obtained in the context of L1 acquisition, a parallel pattern may extend to L2 learners, wherein both the orthographic opacity of the L2 and the typological distance between the L1 and L2 (here, Chinese and English) jointly shape the pace at which visual print becomes coordinated with the spoken-language network.

### 4.5 Limitations

Several limitations of the present study need to be considered. First, the cross-sectional design employed here precludes tracking individual developmental trajectories over time and limits our ability to make inferences about within-subject changes in VWFA organization during L2 literacy acquisition. Longitudinal designs would be essential to directly capture these dynamic developmental processes. Second, while the three groups were carefully matched in age and general cognitive ability as measured by Raven’s Standard Progressive Matrices and differed primarily in their English literacy experience, broader environmental factors such as socioeconomic status (SES) and home literacy environment were not assessed. These factors may contribute to individual differences in L2 literacy acquisition (Noble et al., 2006) and should be systematically considered in future work. Third, the passive viewing task minimized task difficulty differences across groups and was suitable for children with limited English reading experience. However, this approach could not ensure equivalent cognitive load across all groups and categories. Furthermore, recent longitudinal studies have proposed that word representations compete for cortical territory previously involved in processing other visual categories, such as limb images (Kubota et al., 2024; Nordt et al., 2021). As limb stimuli were not included in the present study, this hypothesis could not be directly tested. Future studies incorporating such visual categories would provide valuable insights into the neural competition between word and limb representations across different stages of both L1 and L2 reading acquisition. Finally, future research could benefit from 7T fMRI at millimeter-scale spatial resolution to more precisely characterize how L1 and L2 processing patterns evolve with increasing L2 experience.

### 4.6 Conclusions

In the current study, we examined the systematic modifications and the underlying nature of cortical reorganization in the VWFA during early L2 script acquisition. We found that L2-word selectivity was absent in L2 pre-readers but emerged robustly in beginning readers, while L1-word selectivity remained stable throughout. Representational similarity analysis further revealed that robust L2 word representations within the VWFA emerged only after children began learning to read L2, while L1 word representations remained stable across all three groups. Finally, supervised machine learning classifiers reliably discriminated among the three L2 literacy groups based on L2 activation patterns but not L1 activation patterns, providing no support for the Competing Model’s prediction of competitive neural reallocation away from L1. Together, these findings support the Overlay Model, demonstrating that the VWFA incorporates a new writing system within its existing cortical resources without compromising native-script processing.

## data availability statement

The source data for all figures will be available on Neurovault (https://neurovault.org/) upon acceptance of the paper. Raw data can be made available for academic research upon reasonable request to the corresponding author.

## ethics approval statement

The study was approved by the local ethical committee of University.

