## Supplementary Material for "Learning to read a second language establishes a parallel L2 representation alongside the native one in the VWFA"

### Supplementary Materials

**Supplementary Table S1**

**Within-group correlations between Chinese character recognition and English literacy measures**

| English literacy measure | L2 pre-readers, *r* | L2 beginning readers, *r* | L2 advanced readers, *r* |
|---|---|---|---|
| English spelling | .038 | .367 | .568* |
| English word identification | −.396 | −.167 | .702*** |
| English word attack | −.144 | .005 | .123 |
| English phonological awareness | .115 | .046 | .152 |

Note. Values are Pearson's correlation coefficients between Chinese character recognition and each English literacy measure. *$p < .05$, *** $p < .001$.

**Supplementary Table S2**

**Regions showing significant activations for each category-selective contrast across all participants (voxel-level *p* < .001, uncorrected; cluster-level *p* < .05, FEW-corrected).**

| Region | MNI coordinates | | | No. of voxels in cluster | Cluster-level *p* value (FWE) | Voxel-level *p* value (uncorrected) | Z value at local maximum |
|---|---|---|---|---|---|---|---|
| | x | y | z | | | | |
| **Chinese words > Faces Houses** | | | | | | | |
| Left Precentral Gyrus | −48 | 3 | 48 | 201 | < 0.001 | < 0.001 | 6.67 |
| Left Precentral Gyrus | −45 | 0 | 36 | | | < 0.001 | 5.66 |
| Left Middle Frontal Gyrus | −54 | 15 | 36 | | | < 0.001 | 5.57 |
| Left Middle Temporal Gyrus | −54 | −30 | 0 | 133 | < 0.001 | < 0.001 | 6.57 |
| Left Inferior Temporal Gyrus | −51 | −51 | −15 | 36 | < 0.001 | < 0.001 | 5.80 |
| Left Supplementary Motor Area | −6 | 15 | 54 | 16 | 0.001 | < 0.001 | 5.13 |
| Left Supplementary Motor Area | −6 | 9 | 60 | | | < 0.001 | 5.04 |
| Right Superior Temporal Gyrus | 54 | −27 | 0 | 9 | 0.004 | < 0.001 | 5.13 |
| Left Inferior Frontal Gyrus | −48 | 27 | 3 | 12 | 0.002 | < 0.001 | 4.98 |
| Left Superior Temporal Pole | −54 | 12 | −6 | 3 | 0.014 | < 0.001 | 4.73 |
| Left Superior Temporal Gyrus | −54 | −42 | 21 | 3 | 0.014 | < 0.001 | 4.60 |
| **English words > Faces Houses** | | | | | | | |
| Left Superior Frontal Gyrus | −24 | 0 | 42 | 2 | 0.017 | < 0.001 | 5.01 |
| Left Inferior Frontal Gyrus, Opercular Part | −45 | 3 | 24 | 8 | 0.003 | < 0.001 | 4.96 |
| Left Inferior Temporal Gyrus | −48 | −57 | −15 | 6 | 0.005 | < 0.001 | 4.91 |
| Left Precentral Gyrus | −48 | 0 | 39 | 10 | 0.002 | < 0.001 | 4.87 |
| Left Inferior Parietal Lobule | −42 | −42 | 42 | 2 | 0.017 | < 0.001 | 4.70 |
| **Faces > Others** | | | | | | | |
| Right Cuneus | 18 | −102 | 9 | 90 | < 0.001 | < 0.001 | Inf |
| Right Middle Occipital Gyrus | 30 | −90 | 6 | | | < 0.001 | 5.74 |
| Left Superior Occipital Gyrus | −12 | −105 | 9 | 48 | < 0.001 | < 0.001 | 7.24 |
| Right Fusiform Gyrus | 42 | −51 | −21 | 139 | < 0.001 | < 0.001 | 6.97 |
| Right Inferior Occipital Gyrus | 42 | −78 | −12 | | | < 0.001 | 6.43 |
| Right Fusiform Gyrus | 42 | −60 | −15 | | | < 0.001 | 6.02 |
| Right Hippocampus | 21 | −6 | −18 | 19 | < 0.001 | < 0.001 | 5.68 |
| Left Fusiform Gyrus | −39 | −54 | −18 | 5 | 0.007 | < 0.001 | 5.00 |
| Right Middle Temporal Gyrus | 45 | −60 | 6 | 16 | 0.001 | < 0.001 | 4.99 |
| Left Hippocampus | −18 | −6 | −21 | 3 | 0.013 | < 0.001 | 4.98 |
| Right Parahippocampal Gyrus | 33 | −3 | −30 | 2 | 0.018 | < 0.001 | 4.90 |
| Right Superior Temporal Pole | 30 | 3 | −24 | 3 | 0.013 | < 0.001 | 4.76 |
| Right Inferior Occipital Gyrus | 27 | −93 | −6 | 1 | 0.026 | < 0.001 | 4.66 |
| **Houses > Others** | | | | | | | |
| Left Fusiform Gyrus | −30 | −48 | −9 | 1028 | < 0.001 | < 0.001 | Inf |
| Left Middle Occipital Gyrus | −15 | −102 | 9 | | | < 0.001 | Inf |
| Left Middle Occipital Gyrus | −12 | −99 | 0 | | | < 0.001 | Inf |
| Right Fusiform Gyrus | 30 | −48 | −12 | 1168 | < 0.001 | < 0.001 | Inf |
| Right Fusiform Gyrus | 33 | −69 | −15 | | | < 0.001 | Inf |
| Right Cuneus | 18 | −96 | 12 | | | < 0.001 | Inf |

| | | | | | | | |
|---|---|---|---|---|---|---|---|
| Right Calcarine Fissure and Surrounding Cortex | 21 | −51 | 12 | 7 | 0.005 | < 0.001 | 5.15 |

**Supplementary Table S3**

**Regions showing significant activations for each category-selective contrast in each group (voxel-level *p* < .001, uncorrected; cluster-level *p* < .05, uncorrected).**

| Region | MNI coordinates | | | No. of voxels in cluster | Cluster-level *p* value (uncorrected) | Voxel-level *p* value (uncorrected) | Z value at local maximum |
|---|---|---|---|---|---|---|---|
| | x | y | z | | | | |
| **L2 Pre-readers** | | | | | | | |
| **Chinese words > Faces Houses** | | | | | | | |
| Left Supplementary Motor Area | −3 | 6 | 60 | 53 | 0.025 | < 0.001 | 3.95 |
| Left Inferior Temporal Gyrus | −51 | −48 | −15 | 70 | 0.012 | < 0.001 | 3.76 |
| Left Middle Temporal Gyrus | −51 | −39 | −3 | | | < 0.001 | 3.72 |
| Left Middle Temporal Gyrus | −54 | −48 | −3 | | | < 0.001 | 3.59 |
| **English words > Faces Houses** | | | | | | | |
| No significant clusters. | | | | | | | |
| **Faces > Others** | | | | | | | |
| Right Inferior Occipital Gyrus | 39 | −78 | −12 | 212 | < 0.001 | < 0.001 | 4.86 |
| Right Inferior Occipital Gyrus | 39 | −66 | −12 | | | < 0.001 | 4.42 |
| Right Fusiform Gyrus | 42 | −51 | −21 | | | < 0.001 | 5.25 |
| **Houses > Others** | | | | | | | |
| Left Lingual Gyrus | −30 | −48 | −6 | 700 | < 0.001 | < 0.001 | 5.59 |
| Left Middle Occipital Gyrus | −21 | −102 | 9 | | | < 0.001 | 5.35 |
| Left Middle Occipital Gyrus | −30 | −96 | 12 | | | < 0.001 | 5.33 |
| Right Fusiform Gyrus | 33 | −69 | −15 | 737 | < 0.001 | < 0.001 | 4.89 |
| Right Fusiform Gyrus | 33 | −45 | −9 | | | < 0.001 | 4.76 |
| Right Fusiform Gyrus | 27 | −45 | −15 | | | < 0.001 | 4.73 |
| **L2 Beginning readers** | | | | | | | |
| **Chinese words > Faces Houses** | | | | | | | |
| Left Inferior Frontal Gyrus, Opercular Part | −48 | 12 | 0 | 379 | < 0.001 | < 0.001 | 4.88 |
| Left Inferior Temporal Gyrus | −42 | −51 | −6 | | | < 0.001 | 4.41 |
| Left Superior Temporal Gyrus | −51 | −6 | −6 | | | < 0.001 | 4.36 |
| Left Precentral Gyrus | −51 | 0 | 42 | 199 | < 0.001 | < 0.001 | 4.54 |
| Left Precentral Gyrus | −45 | 3 | 30 | | | < 0.001 | 4.11 |
| Left Precentral Gyrus | −51 | 12 | 33 | | | < 0.001 | 3.91 |
| Left Supplementary Motor Area | −27 | −48 | 39 | 42 | 0.013 | < 0.001 | 3.99 |
| Left Supplementary Motor Area | −39 | −45 | 39 | | | < 0.001 | 3.44 |
| **English words > Faces Houses** | | | | | | | |
| Left Inferior Occipital Gyrus | −45 | −60 | −12 | 43 | 0.019 | < 0.001 | 4.20 |
| Left Precentral Gyrus | −45 | 0 | 33 | 33 | 0.036 | < 0.001 | 3.64 |
| Left Precentral Gyrus | −45 | 3 | 45 | | | < 0.001 | 3.40 |

| | | | | | | | |
|---|---|---|---|---|---|---|---|
| **Faces > Others** | | | | | | | |
| Right Cuneus | 21 | −99 | 9 | 89 | 0.001 | < 0.001 | 4.80 |
| Right Superior Occipital Gyrus | 27 | −90 | 12 | | | < 0.001 | 3.79 |
| Left Superior Occipital Gyrus | −12 | −102 | 9 | 67 | 0.002 | < 0.001 | 4.79 |
| Left Middle Occipital Gyrus | −21 | −99 | 6 | | | < 0.001 | 4.10 |
| Right Fusiform Gyrus | 42 | −51 | −21 | 64 | 0.003 | < 0.001 | 4.42 |
| Right Middle Temporal Gyrus | 42 | −60 | 3 | | | < 0.001 | 4.04 |
| Right Inferior Temporal Gyrus | 48 | −51 | −6 | | | < 0.001 | 3.48 |
| Right Inferior Occipital Gyrus | 39 | −75 | −12 | 32 | 0.024 | < 0.001 | 4.12 |
| Right Inferior Occipital Gyrus | 33 | −72 | −6 | | | < 0.001 | 3.76 |
| Right Inferior Occipital Gyrus | 45 | −75 | −3 | | | < 0.001 | 3.22 |
| Left Lingual Gyrus | −12 | −69 | −3 | 25 | 0.042 | < 0.001 | 4.00 |
| **Houses > Others** | | | | | | | |
| Right Fusiform Gyrus | 30 | −75 | −15 | 776 | < 0.001 | < 0.001 | 5.54 |
| Right Parahippocampal Gyrus | 33 | −42 | −6 | | | < 0.001 | 5.39 |
| Right Parahippocampal Gyrus | 36 | −36 | −15 | | | < 0.001 | 5.08 |
| Left Calcarine Fissure and Surrounding Cortex | −9 | −102 | −3 | 603 | < 0.001 | < 0.001 | 5.18 |
| **L2 Advanced readers** | | | | | | | |
| **Chinese words > Faces Houses** | | | | | | | |
| Left Middle Temporal Gyrus | −57 | −33 | 3 | 92 | 0.003 | < 0.001 | 4.93 |
| Left Middle Temporal Gyrus | −54 | −21 | −3 | | | < 0.001 | 3.55 |
| Left Precentral Gyrus | −36 | 0 | 36 | 159 | < 0.001 | < 0.001 | 4.05 |
| Left Precentral Gyrus | −48 | 6 | 48 | | | < 0.001 | 4.01 |
| Left Inferior Frontal Gyrus, Triangular Part | −54 | 18 | 18 | | | < 0.001 | 3.80 |
| **English words > Faces Houses** | | | | | | | |
| Left Precentral Gyrus | −45 | 3 | 21 | 43 | 0.017 | < 0.001 | 4.04 |
| Left Middle Occipital Gyrus | −27 | −63 | 36 | 29 | 0.043 | < 0.001 | 3.80 |
| **Faces > Others** | | | | | | | |
| Right Calcarine Fissure and Surrounding Cortex | 18 | −102 | 9 | 63 | 0.017 | < 0.001 | 5.49 |
| Right Superior Occipital Gyrus | 27 | −96 | 15 | | | < 0.001 | 3.95 |
| Left Superior Occipital Gyrus | −12 | −105 | 9 | 43 | 0.042 | < 0.001 | 4.81 |
| **Houses > Others** | | | | | | | |
| Right Fusiform Gyrus | 33 | −48 | −15 | 2590 | < 0.001 | < 0.001 | 6.82 |
| Right Fusiform Gyrus | 24 | −42 | −15 | | | < 0.001 | 6.08 |
| Right Fusiform Gyrus | 30 | −66 | −12 | | | < 0.001 | 5.93 |
| Right Precuneus | 21 | −51 | 15 | 63 | 0.005 | < 0.001 | 4.89 |
| Right Calcarine Fissure and Surrounding Cortex | 18 | −48 | 3 | | | < 0.001 | 3.67 |
| Right Rolandic Operculum | 69 | −9 | 12 | 91 | 0.001 | < 0.001 | 3.92 |
| Right Rolandic Operculum | 63 | 6 | 6 | | | < 0.001 | 3.91 |
| Right Postcentral Gyrus | 66 | −3 | 18 | | | < 0.001 | 3.79 |

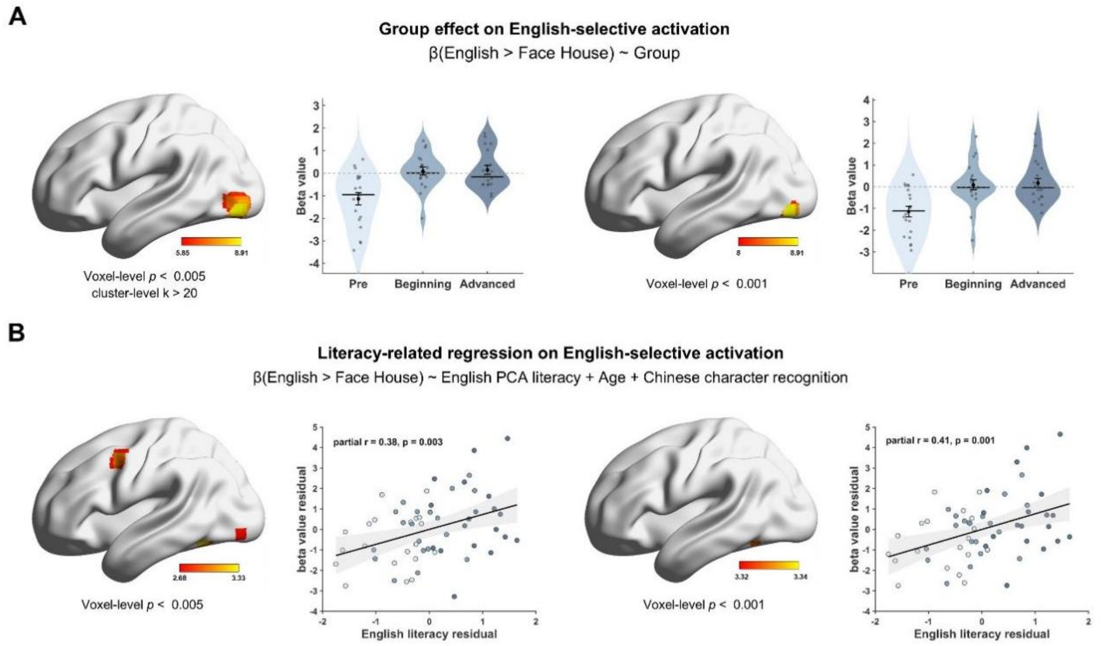


**Supplementary Fig. S1.** Exploratory whole-brain analyses of English-selective activation. (A) Whole-brain one-way ANOVA testing the main effect of group on English-selective activation, defined as English words > the mean of faces and houses. Brain maps are shown at uncorrected voxel-level thresholds of $p < .005$ (left) and $p < .001$ (right) with no additional cluster-extent threshold. Violin plots show mean English-selective contrast estimates extracted from the corresponding suprathreshold clusters for visualization only. (B) Whole-brain regression testing the positive association between English-selective activation and the PCA-based English literacy composite, controlling for age and Chinese character recognition. Brain maps are shown at uncorrected voxel-level thresholds of $p < .005$ (left) and $p < .001$ (right). Scatter plots show partial associations between English literacy and contrast estimates extracted from the suprathreshold cluster near the VWFA, after residualizing both variables with respect to age and Chinese character recognition. No clusters survived whole-brain correction; all extracted-value plots are descriptive.

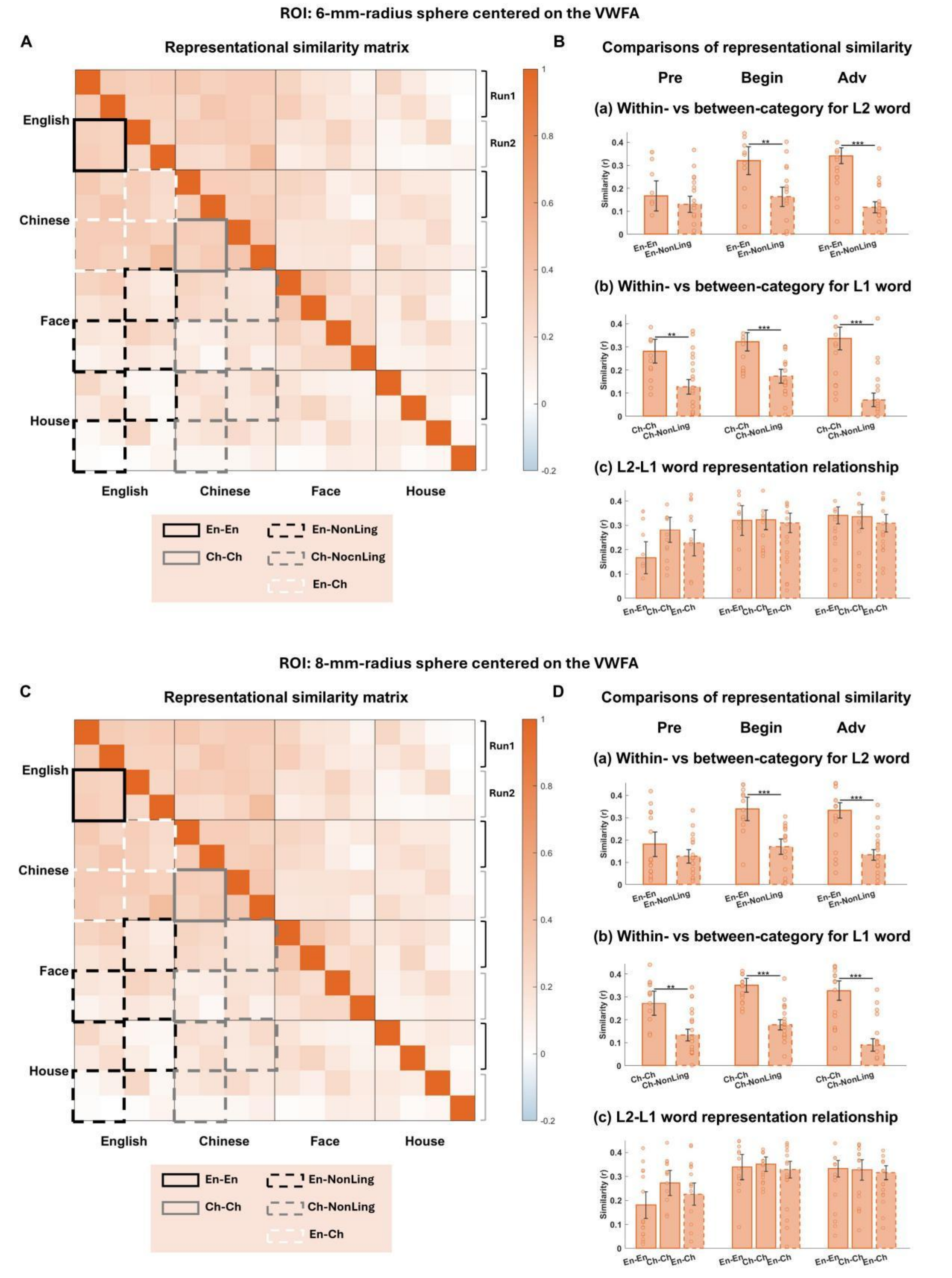


**Supplementary Fig. S2. Representational similarity analyses of word representations in the 6-mm-radius (top panel) and 8-mm-radius spheres (bottom panel) centered on the canonical VWFA coordinates. Left panel:** Representational similarity matrix (RSM) showing the mean, across participants, of the pairwise correlations between spatial activation patterns evoked by four stimulus categories in the 6-mm-radius **(A)** and 8-mm-radius **(C)** spheres. **Right panel:** Comparisons of representational similarity in the 6-mm-radius **(B)** and 8-mm-radius **(D)** spheres centered on the VWFA. Bar plots show mean similarity values for three comparisons: within-English similarity versus English-nonlinguistic similarity, within-Chinese similarity versus Chinese-nonlinguistic similarity, and comparisons among within-English, within-Chinese, and English-Chinese similarity. Results are shown separately for the three groups differing in L2 experience (L2 pre-readers, beginning readers, and advanced readers). Significance levels: *$p_{FDR} < .05$, **$p_{FDR} < .01$, ***$p_{FDR} < .001$ (FDR-corrected paired-sample t-tests).
